# XL-MS and De Novo Protein Design Identified a Common Motif for TREM2 Binding

**DOI:** 10.64898/2026.04.23.720433

**Authors:** Doniesha Perera, Emmanuel Ajiboye, Kalana Pitakatuwana, Sophia Wier, Vincent Duong, Haifan Wu

## Abstract

Apolipoprotein E (*APOE*) and Triggering Receptor Expressed on Myeloid cells 2 (*TREM2*) are the two strongest genetic risk factors of late-onset Alzheimer’s disease. ApoE binds to the low-density lipoprotein receptor (LDLR) to facilitate the uptake of ApoE-lipoprotein particles. TREM2 is a cell surface receptor expressed on microglia in the brain. The activation of TREM2 is essential for microglia to carry out protective functions against AD pathology. Several studies have shown that TREM2 signaling is activated through direct interaction between TREM2 and ApoE. In addition to its important role in AD pathogenesis, the ApoE/TREM2 interaction has been shown to induce immunosuppression of neutrophils within the tumor microenvironment. Despite its clinical importance, a high-resolution molecular understanding of the complex remains elusive. Here, we carried out chemical cross-linking mass spectrometry (XL-MS) analysis of the ApoE3/TREM2_ECD_ complex to identify intra- and inter-protein cross-links, which were used as restraints to guide integrative protein-protein docking. Our data support a binding model in which a helix-loop-helix motif within the ApoE3 hinge and C-terminal region forms a transient hydrophobic pocket that wraps around the hydrophobic tip of the TREM2 ectodomain. This model is further supported by *de novo*-designed mini-protein binders, which show the same binding mode as identified by our XL-MS experiment. These results establish a robust framework for developing mini-protein-based TREM2 agonists.

## INTRODUCTION

Alzheimer’s disease (AD) is a devastating neurodegenerative disorder affecting over 5 million people in the United States. The pathological hallmark of AD is characterized by the deposition of toxic β-amyloid (Aβ) oligomers and fibrils, which initiate a cascade of tau hyperphosphorylation, neurofibrillary tangle formation, and progressive neuronal loss^1–4^. Recent large-scale human genetic studies have highlighted the central role of neuroinflammation and lipid metabolism in AD pathogenesis, identifying apolipoprotein E (*APOE*) and triggering receptor expressed on myeloid cells 2 (*TREM2*) as the two most significant genetic risk factors for late-onset AD^5, 6^. Individuals carrying the *APOEε4* allele face a 10-fold increase in AD risk, while rare variants in the TREM2 gene, such as R47H, confer a 3-fold risk increase.

ApoE is the primary cholesterol and lipid carrier in the central nervous system, mediating the uptake of lipoprotein particles via the low-density lipoprotein receptor (LDLR)^7^. TREM2 is a transmembrane receptor expressed exclusively on microglia in the brain, where it acts as a critical sensor for lipid ligands, apoptotic cells, and Aβ oligomers^8–12^. Activation of TREM2 signaling is essential for microglia to transition into a disease-associated state and mount a protective response against amyloid pathology^13, 14^. Interestingly, ApoE has emerged as a high-affinity putative ligand for TREM2, with *in vitro* studies reporting dissociation constants (K_d_) ranging from 10 to 500 nM^15–20^. Beyond its role in AD, the ApoE-TREM2 signaling axis has recently been implicated in the tumor microenvironment, where it induces immunosuppression of neutrophils, facilitating cancer progression^21^.

Despite the profound clinical implications of the ApoE–TREM2 interaction in both neurodegeneration and oncology, a detailed molecular understanding of the complex remains elusive. Previous studies utilizing site-directed mutagenesis and protein truncation have successfully narrowed the potential binding regions to the ApoE hinge region and the TREM2 CDR loops^18, 20^. Recent computational docking studies have attempted to resolve the complex using the static monomeric NMR structure of ApoE3 (PDB: 2L7B)^19^. However, these studies fail to account for the dynamic nature of ApoE^22^.

In this work, we employed chemical cross-linking coupled with mass spectrometry (XL-MS) to overcome these structural challenges. By capturing intra- and inter-protein distance restraints in solution, we bypassed the limitations of static structural methods. These experimental constraints were integrated with HADDOCK-based docking to generate a 3D model of the TREM2-ApoE3 complex. We also employed *de novo* protein design to further support our binding model.

## RESULTS AND DISCUSSIONS

### XL-MS Analysis of the TREM2/ApoE3 Complex

To map the binding interface between TREM2 and ApoE3, we performed cross-linking mass spectrometry (XL-MS) analysis using the amine-reactive homobifunctional cross-linker BS3. Recombinant TREM2_ECD_ (expressed in *E. coli*) was incubated with commercially available Trx-ApoE3 in the presence of the cross-linker. Analysis of the reaction mixture by SDS-PAGE showed a distinct, albeit faint, band above the Trx-ApoE3 band, corresponding to the expected molecular weight of the heterodimeric TREM2_ECD_/Trx-ApoE3 complex (**Figure 1a**). Additionally, higher-molecular-weight species were detected, suggesting the formation of larger oligomeric species, such as tetramers.

**Figure 1.**
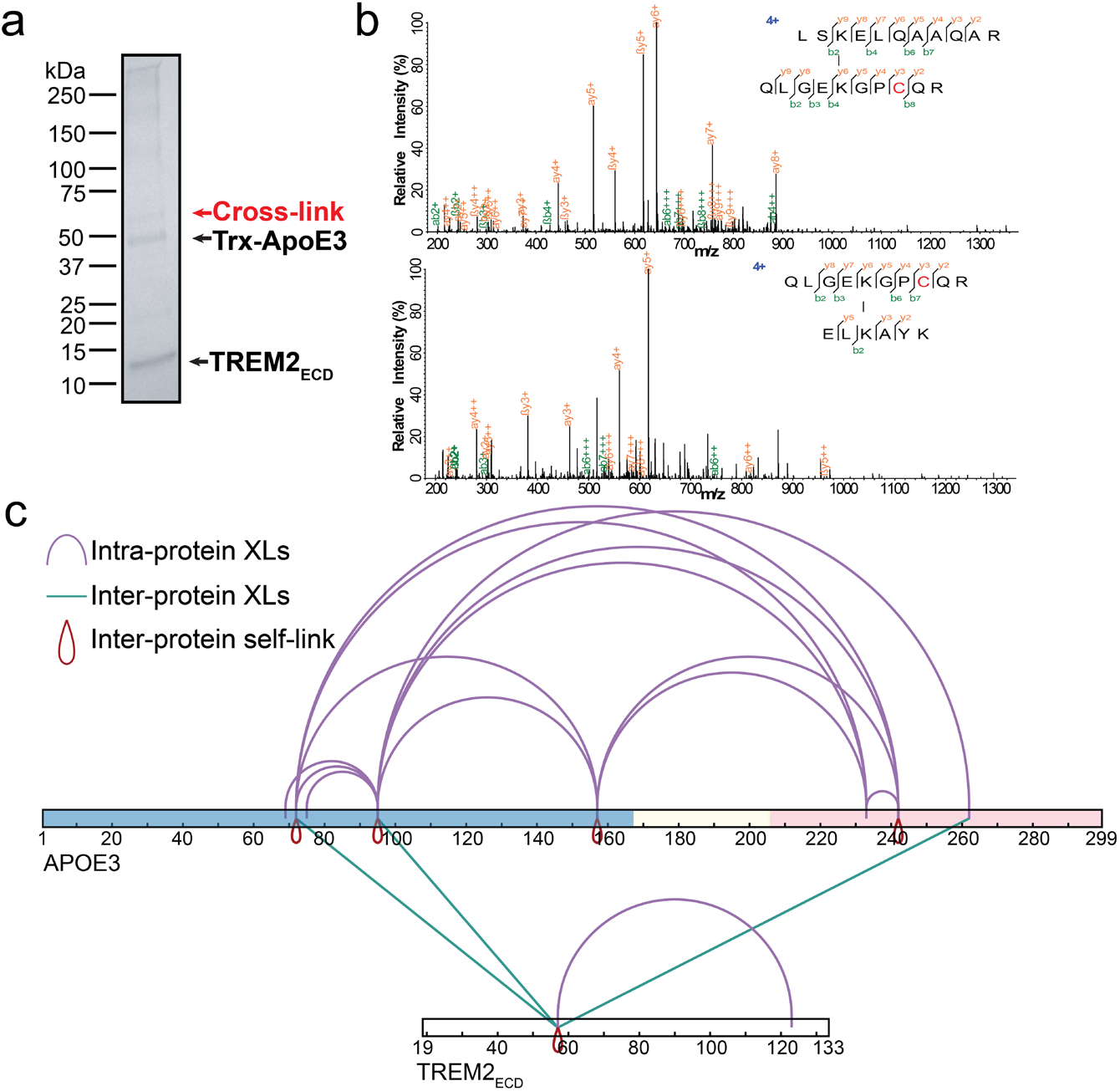
Chemical cross-linking of Trx-ApoE3 and TREM2_ECD_. (a) SDS-PAGE analysis showing successful cross-linking of the heterodimeric TREM2_ECD_/Trx-ApoE3 complex. (b) Representative high-quality MS/MS spectra of inter-protein cross-linked peptides. (c) Bar representation showing intra-protein XLs, inter-protein XLs, and inter-protein self-links. Figure was created using xiNET^23^. ApoE3: light blue (N-terminal region, residues 1-167), light yellow (hinge region, residues 168-205), and pink (C-terminal region, residues 206-299).

The cross-linked mixture was subjected to tryptic digestion and analyzed via LC-MS/MS to identify specific cross-linking sites. To ensure high confidence in our structural constraints, only cross-linked peptides identified across three technical replicates were used for downstream analysis. Several inter-protein cross-links were supported by high-quality MS/MS spectra (**Figure 1b**), confirming the physical proximity of these residues within the TREM2_ECD_/Trx-ApoE3 interface.

The identified cross-links were mapped onto the primary sequences of both proteins (**Dataset1, Figure 1c, and Table S1**). In addition to the 14 intra-protein and 3 inter-protein cross-links, we also detected inter-protein self-links (cross-links between two identical subunits), which agrees with the higher-order oligomerization observed by SDS-PAGE. While the intra-protein constraints provided internal validation for the folding of the individual proteins, the inter-protein cross-links served as important distance restraints for the subsequent protein-protein docking study.

### Conformational Filtering and Docking

Initial data analysis using pLink3 identified one intra-TREM2 cross-link and 13 intra-ApoE3 cross-links. The intra-protein constraint within TREM2 was successfully mapped onto the crystal structure (PDB: 5ELI), yielding a Cβ-Cβ solvent-accessible surface distance (SASD) below the 35 Å maximum threshold for BS3. This confirmed the structural integrity of the recombinant TREM2_ECD_. In contrast, mapping the 13 intra-ApoE3 cross-links onto the representative NMR structure (PDB: 2L7B)^24^ revealed significant structural discrepancies. The majority (11 out of 13) of the identified cross-links exhibited Cβ-Cβ SASDs exceeding 35 Å (**Figure 2a**). Furthermore, several cross-linked lysine residues were buried within the protein core of 2L7B, making them solvent-inaccessible and unable to react with the BS3 cross-linker. These observations indicated that the static 2L7B structure does not accurately represent the solution-state conformation of ApoE3 used in our XL-MS experiment.

**Figure 2.**
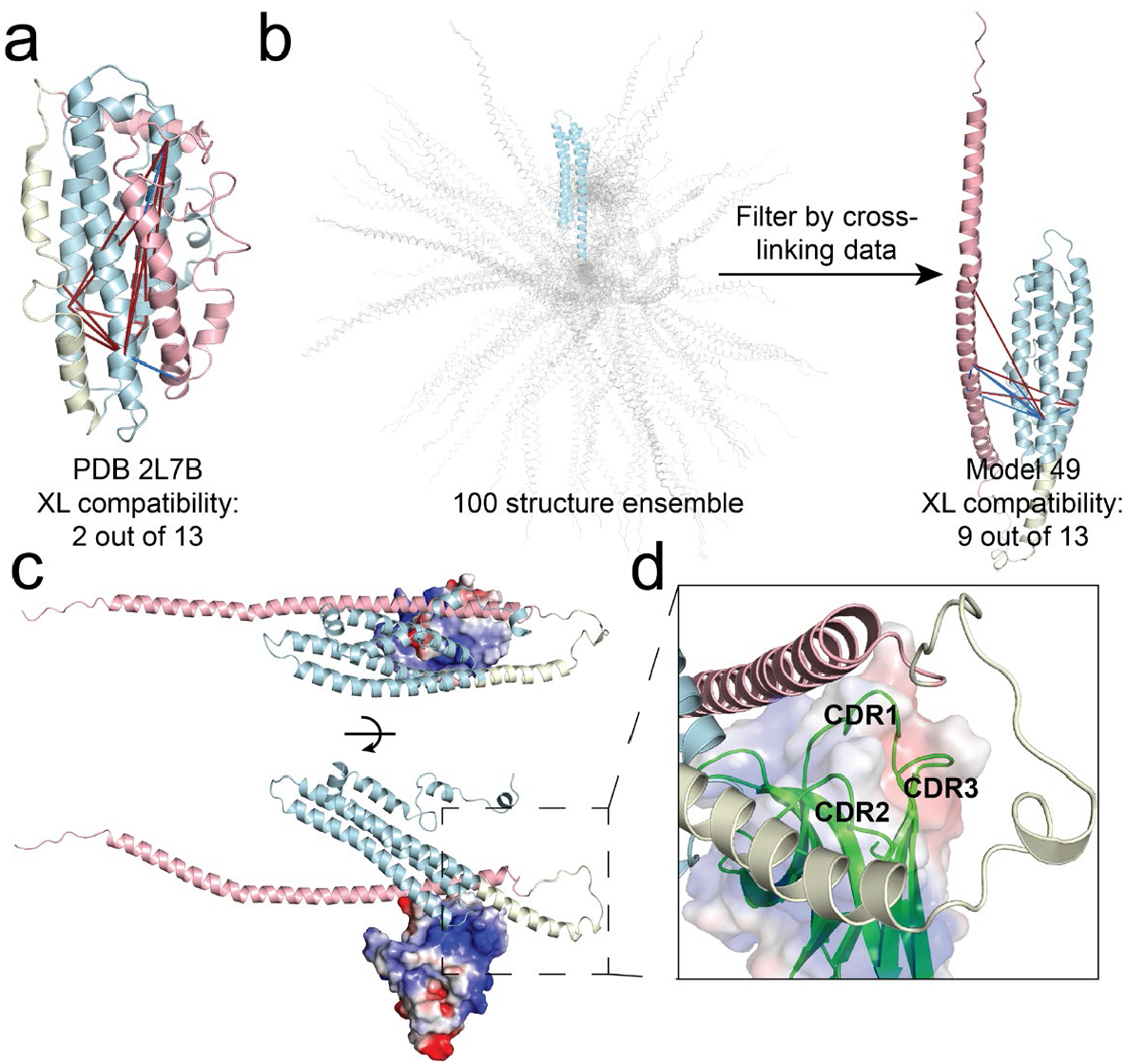
Modeling TREM2/ApoE3 complex using XL-MS data. (a) Mapping intra-ApoE3 cross-links onto the NMR structure (pdb 2L7B). Red line: incompatible XLs with Cβ-Cβ solvent accessible surface distance (SASD) > 35 Å. Blue line: compatible XLs. (b) Filtering the structure ensemble of ApoE3 (Protein Ensemble Database PED07094) by intra-ApoE3 XLs yielded Model 49 with the highest structural compatibility. (c) The best-scoring model of TREM2/ApoE3 complex generated using Haddock. (d) Zoom-in view of the binding interface. ApoE3: light blue (N-terminal region, residues 1-167), light yellow (hinge region, residues 168-205), and pink (C-terminal region, residues 206-299). TREM2_ECD_: CDR1(residue 40-42), CDR2(residue 69-72), and CDR3 (residue 88-91).

To identify a more relevant conformation, we utilized a 100-structure ensemble of ApoE3 from the Protein Ensemble Database (PED07094, **Figure 2b**). These structures were generated via AlphaFlex with IDPConformerGenerator and energy minimized by molecular dynamics simulation^25^. Each structure was filtered based on its compatibility with our experimental XL-MS data (**Table S2**). Model 49 was identified as the top-ranking conformation, exhibiting the highest percentage of satisfaction (8 out of 13 cross-links), and was subsequently used as the template for docking.

In addition to the intra-protein constraints, three high-confidence inter-protein cross-links were identified (**Table S1**). These cross-links, combined with mutagenesis data^18^, were utilized as ambiguous restraints for rigid-body docking using HADDOCK^26, 27^. The docking study yielded 191 structures in six clusters. The lowest-energy model reveals a binding interface on ApoE3 that spans both the C-terminal lipid-binding region and the flexible hinge region (**Figure 2c**). The binding interface on TREM2 is at the hydrophobic tip of the Ig fold. This recognition site involves hydrophobic residues within the complementarity-determining regions (CDRs 1–3). The interface is characterized by a helix-loop-helix motif from ApoE3 that docks onto the hydrophobic tip of TREM2 (**Figure 2d**).

Although the previous docking study^19^ of the TREM2/ApoE3 interface relied on the monomeric NMR structure of ApoE3 (PDB: 2L7B), our cross-linking mass spectrometry (XL-MS) data reveal that this structure represents a conformational state incompatible with TREM2 binding. Instead, we propose an ‘induced fit’ mechanism in which the flexible hinge and C-terminal regions undergo a conformational transition to form a transient hydrophobic pocket. This pocket provides the necessary surface complementarity to accommodate the hydrophobic tip of the TREM2 ectodomain, specifically engaging the CDR1–3 loops. This model provides a potential structural explanation for how TREM2 recognizes ApoE across its diverse lipid-binding states^18^.

### *De novo* design of mini-protein binders

To further support the binding model, we employed *de novo* protein design using BindCraft^28^ and Odesign ([arXiv:2510.22304](https://arxiv.org/abs/2510.22304)). We restricted the design space to mini-proteins of 30–75 residues to ensure feasibility for chemical synthesis and subsequent site-specific modifications. The BindCraft pipeline generated 100 designs, while Odesign yielded 6 refined scaffolds. From this pool, a total of 13 leads—comprising the top 10 BindCraft and 3 Odesign designs—were selected for chemical synthesis and experimental characterization (**Figure 3, Table S3 and S4**). The binding affinities of synthetic mini-proteins were measured using microscale thermophoresis (**Figures 3 and SI**). 8 out of 13 tested designs exhibited measurable binding, with dissociation constants (K_d_) in the nanomolar to single-digit micromolar range. BindCraft8 and Odesign2 are top binders, achieving K_d_ values of 370 and 80 nM, respectively. Remarkably, even though no “hotspot” residues were specified during the design process, these designs converged on the hydrophobic tip of TREM2. They primarily engage the CDR loops, especially the CDR2 loop, resembling the docking architecture proposed in our ApoE3/TREM2 model.

**Figure 3.**
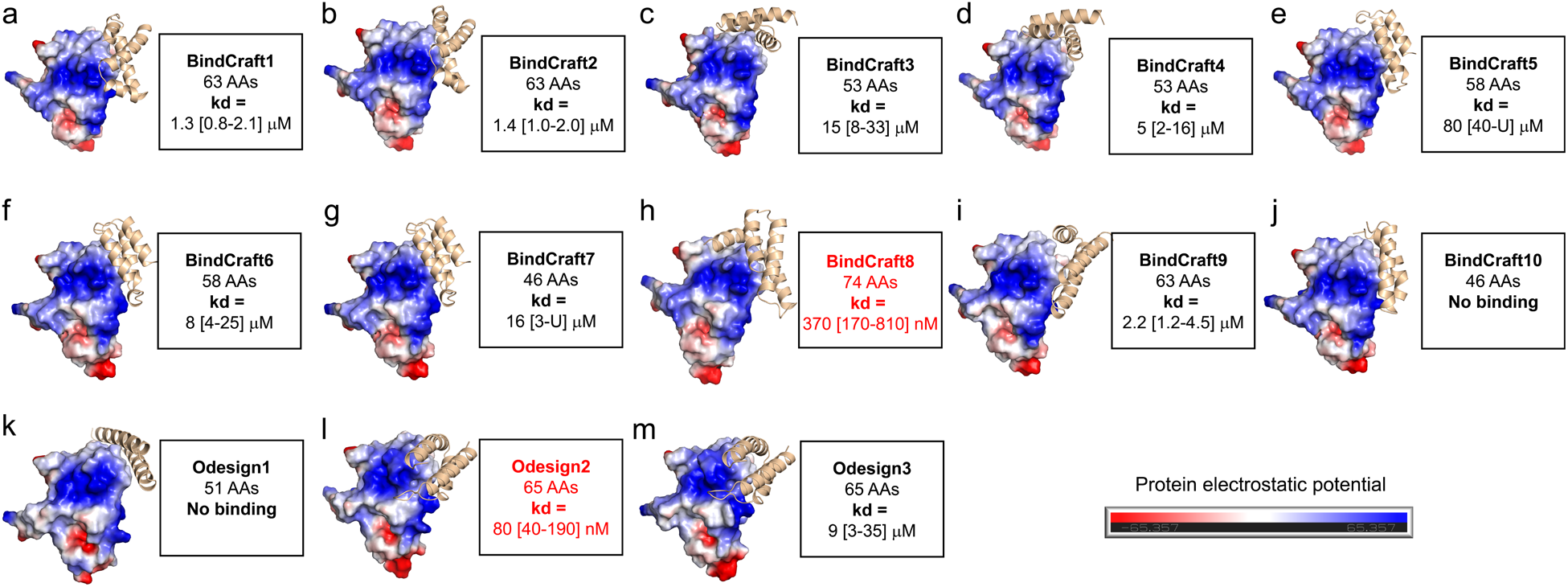
De novo mini-protein binder design. (a-m) Design models of mini-proteins in complex with TREM2_ECD_ (left) and binding affinity determined by microscale thermophoresis (right). Numbers in brackets represent the 68.3% confidence interval calculated by “error-surface projection”^29^.

**Figure 4.**
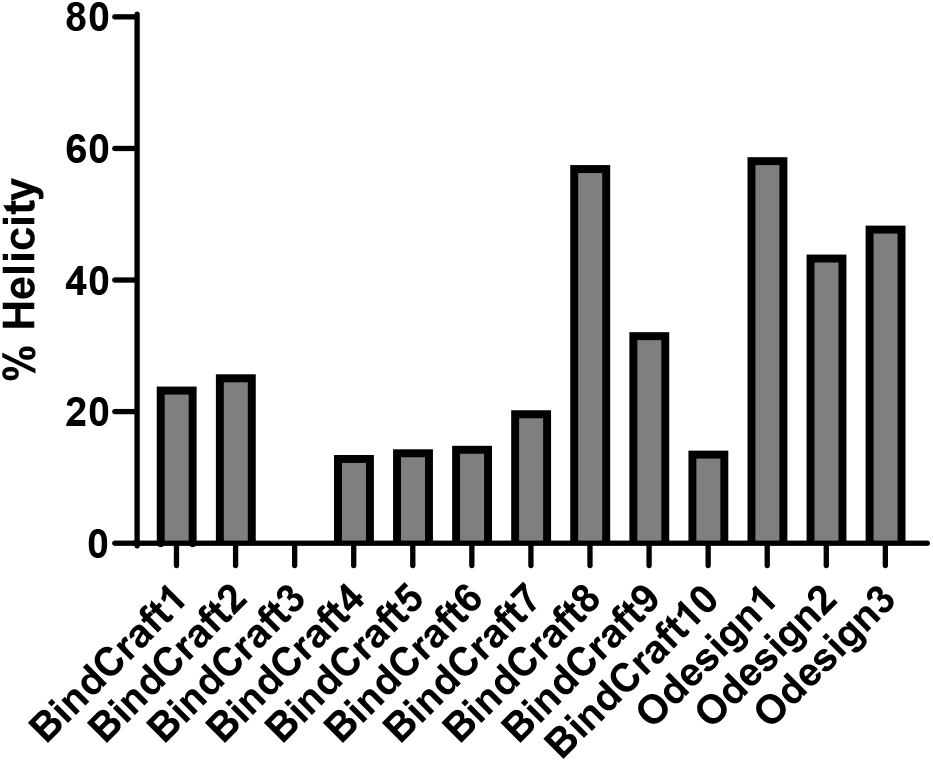
Percentage helicity calculated for each designed mini-protein based on CD spectrum.

To investigate the structural basis for the observed binding affinities, we analyzed the secondary structures of the 13 mini-proteins using circular dichroism (CD) spectroscopy. The two top binders, BindCraft8 and Odesign2, exhibited high helical content, consistent with their designed folds. Conversely, weak and non-binding designs (e.g., BindCraft3–7 and 10) showed significantly lower helicity, suggesting poor structural stability. Surprisingly, Odesign1 and Odesign3, despite displaying high helicity, exhibited weak binding, likely due to suboptimal shape complementarity at the TREM2 interface. Furthermore, smaller designs such as BindCraft7, 10, and Odesign1 failed to show binding, indicating that a threshold interaction surface area is required for stable complex formation. These data suggest that structural pre-organization and sufficient interaction surfaces are critical determinants of potent binding.

## CONCLUSIONS

In summary, we have utilized an integrative XL-MS and computational modeling approach to identify the structural basis of the TREM2-ApoE3 interaction. Our data revealed a helix-loop-helix motif within the ApoE3 hinge and C-terminal region that forms a transient hydrophobic pocket, wrapping around the hydrophobic tip of the TREM2 ectodomain. To support this model, we employed de novo protein design to generate targeted mini-protein binders. Remarkably, without the use of pre-specified hotspots, our top binders converged onto the same hydrophobic tip identified by our XL-MS model, achieving nanomolar binding potency. To our knowledge, this represents the first successful *de novo* design of a targeted mini-protein for the TREM2 ectodomain and provides a robust foundation for developing mini-protein based TREM2 agonists.

## METHODS

### Materials

All Fmoc protected amino acids, HCTU, DIEA, MPAA, TFA, 4-methylpiperidine, and TIPS were purchased from Chemimpex Inc. HPLC grade TFA was purchased from VWR. 2-chlorotrityl resin and Rink resin were purchased from Chempep Inc. TECP was purchased from TCI Chemicals. HPLC grade acetonitrile was purchased from VWR. All other reagents were purchased from Fisher Scientific.

### Preparation of hydrazine resin

The hydrazine resin was freshly prepared before peptide synthesis. Briefly, 200 mg 2-chlorotrityl resin was swelled in 2 mL DMF for 15 min. A mixture of DIEA (125 μL) and 10% hydrazine hydrate in DMF (380 μL) was added dropwise to the resin, and the suspension was shaken at room temperature for 2 h. 2 mL MeOH was then added to the suspension, which was shaken for additional 15 min. Finally, the resin was washed with DMF (3 mL, x4). The C-terminal amino acid was introduced to the hydrazine resin by treating resin with Fmoc-AA (4 eq.), DIEA (16 eq.) and HCTU (3.8 eq) in 2 mL DMF for 1 h at room temperature. The coupling reaction was repeated.

### Solid-phase peptide synthesis

#### Synthesis of peptides with C-terminal hydrazide peptides

Solid-phase peptide synthesis (SPPS) was conducted on a Purepep Chorus peptide synthesizer with the hydrazine resin (0.1 mmol scale). A typical SPPS cycle includes Fmoc deprotection (3 mL 20% 4-methylpiperidine/DMF, 10-min shaking at rt, repeated), DMF wash (4 mL, x3), double coupling (4 eq. Fmoc-AA, 3.8 eq. HCTU, 8 eq. DIEA in 4 mL DMF, 10-min shaking at 75 °C), and another DMF washing step (4 mL, x3). After SPPS cycles were completed, a final Fmoc deprotection step was conducted.

#### Synthesis of peptides with C-terminal amide

Solid-phase peptide synthesis (SPPS) was conducted on a Purepep Chorus peptide synthesizer with the Rink amide resin (0.1 mmol scale). A typical SPPS cycle includes Fmoc deprotection (3 mL 20% 4-methylpiperidine/DMF, 10-min shaking at rt, repeated), DMF wash (4 mL, x3), double coupling (4 eq. Fmoc-AA, 3.8 eq. HCTU, 8 eq. DIEA in 4 mL DMF, 10-min shaking at 75 °C), and another DMF washing step (4 mL, x3). After SPPS cycles were completed, a final Fmoc deprotection step was conducted.

#### Peptide cleavage

The resin was washed with DMF (4 mL, x3) and DCM (4 mL, x5) and dried with N_2_ flow for 20 min. Peptide cleavage was done using 5 mL TFA cleavage cocktail (TFA:TIPS:H_2_O:DTT, 94:2:2:2) for 3 h. The resin slurry was filtered, and the filtrate was added dropwise to 40 mL cold diethyl ether (pre-chilled on dry ice). White precipitate was collected by centrifugation and washed with 40 mL cold ether. Crude peptide was obtained after air drying.

#### General procedure of HPLC purification

The crude peptide was dissolved in 10 mL 6 M GnHCl and filtered through a 0.45 μm syringe filter. Crude material was purified by reverse-phase HPLC using a C4 preparative column (Higgins Analytical, Inc.) and solvent A (water 0.1% TFA)/B (ACN, 0.1% TFA) as the mobile phase.

### Native chemical ligation

The peptide hydrazide was dissolved in an activation buffer (6 M GnHCl, 100 mM sodium phosphate, pH 3.0) to 2 mM. The solution was sonicated for 10 min and chilled on salt/ice bath for 10 min. The activation of hydrazide was done by adding freshly prepared NaNO_2_ solution (0.3 M in the activation buffer) to reach a final concentration of 30 mM. The mixture was incubated on salt/ice bath for 20 min. Then, an equal volume of a freshly prepared MPAA solution (0.2 M MPAA in 6 M GnHCl, 100 mM sodium phosphate, pH 7.5) was added to the mixture. The resulting solution was allowed to warm up to RT and added to the N-cys peptide. After 10-min sonication, the final pH was adjusted to 6.8-7.0 by addition of 2 M NaOH (increment of 2 μL). The solution was kept at RT for overnight. Upon completion, an equal volume of TCEP solution (0.1 M in GnHCl, pH 7) was added to the reaction mixture, and the solution was incubated for 10 min. The solution was acidified by mixing with TCEP solution (0.1 M in GnHCl, pH 2). After centrifugation at 14,000 g for 10 min, the supernatant was collected and purified by reverse-phase HPLC.

### Expression, refolding, and purification of TREM2 ectodomain

The expression, refolding, and purification of TREM2 ectodomain were carried out following a previously published procedure^30^.

### Cross-linking mass spectrometry (XL-MS)

#### Protein cross-linking

20 μM Trx-ApoE3 (Sino Biological 10817-H30E) and 20 μM TREM2 ectodomain were incubated for 1 h at 4°C. The complex was cross-linked by 1 mM BS3 in a cross-linking buffer (20 mM HEPES, 150 mM NaCl, pH 7.5) at rt. The reaction was quenched by adding 20 mM NH_4_HCO_3_ and incubated at rt for 15 min. The reaction mixture was then treated with 6 volumes of cold acetone (−20 °C) and incubated at −20 °C for 1 hour to precipitate the protein.

#### Mass spec sample preparation

The precipitated protein pellets were air dried and resuspended in 8 M urea, 100 mM Tris, pH 8.5, then reduced with 5 mM DTT for 30 min at 37 °C and alkylated with 14 mM iodoacetamide for 15 min in the dark at RT. After reduction and alkylation, samples were diluted to 1 M urea with 100 mM Tris (pH 8.5) and digested with Trypsin/Lys-C Mix (Promega, protein: enzyme ratio 25:1) at 37 °C for overnight. Digestion was stopped by adding formic acid to 5% final concentration, and digested peptides were desalted with C18 Zip tips.

#### LC-MS/MS analysis

Tryptic peptides were separated by reverse phase XSelect CSH C18 2.5 um resin (Waters) on an in-line 150 x 0.075 mm column using an UltiMate 3000 RSLCnano system (Thermo). Peptides were eluted using a 60 min gradient from 98:2 to 65:35 buffer A:B ratio. Eluted peptides were ionized by electrospray (2.4 kV) followed by mass spectrometric analysis on an Orbitrap Eclipse Tribrid mass spectrometer (Thermo). MS data were acquired using the FTMS analyzer in profile mode at a resolution of 120,000 over a range of 375 to 1400 m/z with advanced peak determination. Following HCD activation, MS/MS data were acquired using the FTMS analyzer in profile mode at a resolution of 15,000 over a range of 150 to 2000 m/z with a stepped collision energy of 27-33%. Buffer A = 0.1% formic acid, 0.5% acetonitrile. Buffer B = 0.1% formic acid, 99.9% acetonitrile.

#### XL-MS data analysis

Raw data were analyzed by the pLink 3 software (version 3.0.17)^31^ with the following search parameters: linker set to BS3, enzyme set to trypsin_P, precursor mass accuracy at 10 ppm, fragment mass accuracy at 20 ppm, fixed modification set to carbamidomethyl, variable modification set to oxidation[M], peptide mass ranged from 600 to 6,000 Da, and the minimum and maximum peptide length 6 and 60. Cross-links identified in all three technical replicates were chosen for visualization by xiNET^23^ and further analysis of solvent accessible surface distance (SASD) by Xwalk^32^.

### Protein structure modeling

#### Structural Ensemble Filtering via XL-MS SASD compatibility

An ensemble of 100 ApoE3 structures (https://proteinensemble.org/, PED07094)^25, 33^ was analyzed to identify conformations consistent with experimental cross-linking data. Intra-ApoE cross-links were mapped onto each state of the ensemble. For every structure, the Solvent Accessible Surface Distance (SASD) was calculated for all identified cross-linked pairs. A “compatibility score” was assigned based on the percentage of experimental links that satisfied the SASD threshold (35 Å). The structure exhibiting the highest compatibility was selected as the representative template for subsequent docking and binding mode analysis.

#### Protein docking using HADDOCK

Protein-protein docking was performed using HADDOCK 2.4 webserver^27^ (https://wenmr.science.uu.nl/haddock2.4/) following the procedures and settings described in this tutorial (https://www.bonvinlab.org/education/HADDOCK-Xlinks/). The docking was guided by both experimental cross-linking constraints and mutagenesis data^18^. Model 1 was defined as the validated ApoE3 structure (Model 49 from the filtered PED07094 ensemble), and Model 2 was the crystal structure of the TREM2 ectodomain (PDB: 5ELI, chain B). Residues 191-238 of ApoE3 and residues (41, 44, 69, 70, 74) of TREM2 were set as active residues, and residues 54-59 of TREM2 were assigned as semi-flexible residues. Inter-protein distance restraints derived from XL-MS were incorporated as unambiguous restraints, ensuring the final models were consistent with the experimental spatial topography.

### Protein design

#### BindCraft

Protein design was done using the BindCraft colab notebook (https://colab.research.google.com/github/martinpacesa/BindCraft/blob/main/notebooks/BindCraft.ipynb)^28^. Chain B of pdb 5ELI was used as the target without hotspot selection, and design length was set between 30 to 75 residues. 100 final designs were required. The top 10 designs were selected for chemical synthesis and validation.

#### Odesign

Protein design was done using the Odesign webserver (https://odesign.lglab.ac.cn). Chain B of pdb 5ELI was used as the target without hotspot selection, and design length was set between 20 to 50 residues (for Odesign1,5,6) and 65 residues (for Odesign2,3,4). The top 3 designs based on Binder PTM (>0.8) were selected for chemical synthesis and validation.

### Microscale thermophoresis

His-tagged TREM2(19-174) protein (Sino Biological 11084-H08H-UE) was fluorescently labeled using RED-tris-NTA labeling kit (NanoTemper Technologies). The concentration of labeled TREM2 was kept constant at 10 nM in the final mixture, while the unlabeled mini-protein binder was prepared in a 16-step two-fold serial dilution ranging from 20 μM to 0.3 nM. Samples were loaded into Monolith NT.115 premium coated capillaries. Measurements were performed on a Monolith NT.115 instrument at 25°C in a binding buffer containing 20 mM HEPES pH 7.4, 150 mM NaCl, and 0.05% Tween-20. The MST power was set to medium, and the LED excitation power was set to auto detect. Data analysis was performed using PALMIST^29^.

### CD spectroscopy

Designed mini-proteins (0.2 mg/mL) were dissolved in 10 mM Na phosphate, 100 mM NaF, pH 7.4. Analysis was done on a Jasco 810 CD spectrometer using a 1 mm cuvette at rt. The percent helicity was calculated using the BeStSel webserver (https://bestsel.elte.hu/index.php)^34^.

## ASSOCIATED CONTENT

### Supporting Information

The Supporting Information is available free of charge.

HPLC traces and MST data analysis of mini-proteins; cross-link list; mini-protein sequences and scores.

## AUTHOR INFORMATION

### Author Contributions

D.P., E.A., and H.W. designed research; D.P., E.A., K.P., S.W., V.D., and H.W. performed experiments; D.P. and H.W. analyzed data and wrote the paper.

## ACKNOWLEDGMENTS

Research was supported by the National Institute on Aging of the National Institutes of Health under award number R15AG080493.

We thank Drs. Matthew Hart and Nataliya Smith in Small Molecule and CRISPR High-Throughput Screening Facility (HTSF) at University of Oklahoma Medical Center for help with MST analysis and Dr. Sam Mackintosh in IDeA National Resource for Quantitative Proteomics (R24GM137786) for help with XL-MS.

## ABBREVIATIONS

AD: Alzheimer’s disease
APOE: apolipoprotein E
TREM2: Triggering receptor expressed on myeloid cells 2
Aβ: Amyloid β
SPPS: Solid Phase Peptide Synthesis
CD: Circular Dichroism
MST: Microscale thermophoresis
XL-MS: cross-linking mass spectrometry
NCL: Native Chemical Ligation
HPLC: High Performance Liquid Chromatography
ECD: Ectodomain
Ig fold: Immunoglobin fold
AA: Amino Acid
HCTU: *O*-(1*H*-6-Chlorobenzotriazole-1-yl)-1,1,3,3-tetramethyluronium hexafluorophosphate
DIEA: N,N-Diisopropylethylamine
MPAA: 4-Mercaptophenylacetic acid
TFA: Trifluoroacetic Acid
TIPS: Triisopropyl silane
DMF: Dimethylformamide
TCEP: tris(2-carboxyethyl)phosphine
RT: Room Temperature
GndCl: Guanidinium chloride
DTT: Dithiothreitol
Tris-HCl: Tris Hydrochloride
HEPES: 2-[4-(2-Hydroxyethyl)piperazin-1-yl]ethane-1-sulfonic acid.

