## Supplementary material for "XL-MS and De Novo Protein Design Identified a Common Motif for TREM2 Binding": SI

**Table S1. A list of cross-links**

| Protein1 | LinkPos1 | Protein2 | LinkPos2 | Link_Type |
|---|---|---|---|---|
| TREM2 | 57 | ApoE3 | 72 | Inter-chain |
| TREM2 | 57 | ApoE3 | 95 | Inter-chain |
| TREM2 | 57 | ApoE3 | 262 | Inter-chain |
| ApoE3 | 72 | ApoE3 | 233 | Intra-chain |
| ApoE3 | 233 | ApoE3 | 242 | Intra-chain |
| ApoE3 | 157 | ApoE3 | 233 | Intra-chain |
| ApoE3 | 95 | ApoE3 | 233 | Intra-chain |
| ApoE3 | 72 | ApoE3 | 157 | Intra-chain |
| ApoE3 | 95 | ApoE3 | 262 | Intra-chain |
| ApoE3 | 72 | ApoE3 | 95 | Intra-chain |
| ApoE3 | 75 | ApoE3 | 95 | Intra-chain |
| ApoE3 | 72 | ApoE3 | 72 | Intra-chain |
| ApoE3 | 95 | ApoE3 | 242 | Intra-chain |
| ApoE3 | 95 | ApoE3 | 157 | Intra-chain |
| ApoE3 | 72 | ApoE3 | 242 | Intra-chain |
| ApoE3 | 157 | ApoE3 | 242 | Intra-chain |
| ApoE3 | 69 | ApoE3 | 95 | Intra-chain |
| TREM2 | 57 | TREM2 | 123 | Intra-chain |
| ApoE3 | 95 | ApoE3 | 95 | Intra-chain |
| TREM2 | 57 | TREM2 | 57 | Intra-chain |
| ApoE3 | 242 | ApoE3 | 242 | Intra-chain |
| ApoE3 | 157 | ApoE3 | 157 | Intra-chain |

**Table S2. Compatibility scores of 100 ApoE3 structures**

| Rank | Model | Compatibility score | Valid_XLs | Total_XLs |
|---|---|---|---|---|
| 1 | 49 | 61.54 | 8 | 13 |
| 2 | 96 | 61.54 | 8 | 13 |
| 3 | 35 | 53.85 | 7 | 13 |
| 4 | 6 | 46.15 | 6 | 13 |
| 5 | 51 | 46.15 | 6 | 13 |
| 6 | 76 | 46.15 | 6 | 13 |
| 7 | 91 | 46.15 | 6 | 13 |
| 8 | 2 | 38.46 | 5 | 13 |
| 9 | 3 | 38.46 | 5 | 13 |
| 10 | 4 | 38.46 | 5 | 13 |
| 11 | 5 | 38.46 | 5 | 13 |
| 12 | 7 | 38.46 | 5 | 13 |
| 13 | 8 | 38.46 | 5 | 13 |

| | | | | |
|---|---|---|---|---|
| 14 | 9 | 38.46 | 5 | 13 |
| 15 | 10 | 38.46 | 5 | 13 |
| 16 | 11 | 38.46 | 5 | 13 |
| 17 | 12 | 38.46 | 5 | 13 |
| 18 | 13 | 38.46 | 5 | 13 |
| 19 | 14 | 38.46 | 5 | 13 |
| 20 | 15 | 38.46 | 5 | 13 |
| 21 | 17 | 38.46 | 5 | 13 |
| 22 | 18 | 38.46 | 5 | 13 |
| 23 | 19 | 38.46 | 5 | 13 |
| 24 | 20 | 38.46 | 5 | 13 |
| 25 | 21 | 38.46 | 5 | 13 |
| 26 | 24 | 38.46 | 5 | 13 |
| 27 | 28 | 38.46 | 5 | 13 |
| 28 | 29 | 38.46 | 5 | 13 |
| 29 | 31 | 38.46 | 5 | 13 |
| 30 | 32 | 38.46 | 5 | 13 |
| 31 | 33 | 38.46 | 5 | 13 |
| 32 | 34 | 38.46 | 5 | 13 |
| 33 | 36 | 38.46 | 5 | 13 |
| 34 | 37 | 38.46 | 5 | 13 |
| 35 | 38 | 38.46 | 5 | 13 |
| 36 | 39 | 38.46 | 5 | 13 |
| 37 | 40 | 38.46 | 5 | 13 |
| 38 | 42 | 38.46 | 5 | 13 |
| 39 | 44 | 38.46 | 5 | 13 |
| 40 | 45 | 38.46 | 5 | 13 |
| 41 | 48 | 38.46 | 5 | 13 |
| 42 | 50 | 38.46 | 5 | 13 |
| 43 | 52 | 38.46 | 5 | 13 |
| 44 | 53 | 38.46 | 5 | 13 |
| 45 | 55 | 38.46 | 5 | 13 |
| 46 | 57 | 38.46 | 5 | 13 |
| 47 | 58 | 38.46 | 5 | 13 |
| 48 | 59 | 38.46 | 5 | 13 |
| 49 | 60 | 38.46 | 5 | 13 |
| 50 | 61 | 38.46 | 5 | 13 |
| 51 | 62 | 38.46 | 5 | 13 |
| 52 | 63 | 38.46 | 5 | 13 |
| 53 | 67 | 38.46 | 5 | 13 |
| 54 | 68 | 38.46 | 5 | 13 |
| 55 | 69 | 38.46 | 5 | 13 |
| 56 | 70 | 38.46 | 5 | 13 |

| | | | | |
|---|---|---|---|---|
| 57 | 71 | 38.46 | 5 | 13 |
| 58 | 72 | 38.46 | 5 | 13 |
| 59 | 74 | 38.46 | 5 | 13 |
| 60 | 78 | 38.46 | 5 | 13 |
| 61 | 80 | 38.46 | 5 | 13 |
| 62 | 84 | 38.46 | 5 | 13 |
| 63 | 85 | 38.46 | 5 | 13 |
| 64 | 88 | 38.46 | 5 | 13 |
| 65 | 89 | 38.46 | 5 | 13 |
| 66 | 92 | 38.46 | 5 | 13 |
| 67 | 93 | 38.46 | 5 | 13 |
| 68 | 95 | 38.46 | 5 | 13 |
| 69 | 97 | 38.46 | 5 | 13 |
| 70 | 98 | 38.46 | 5 | 13 |
| 71 | 99 | 38.46 | 5 | 13 |
| 72 | 1 | 30.77 | 4 | 13 |
| 73 | 16 | 30.77 | 4 | 13 |
| 74 | 22 | 30.77 | 4 | 13 |
| 75 | 23 | 30.77 | 4 | 13 |
| 76 | 25 | 30.77 | 4 | 13 |
| 77 | 26 | 30.77 | 4 | 13 |
| 78 | 27 | 30.77 | 4 | 13 |
| 79 | 30 | 30.77 | 4 | 13 |
| 80 | 43 | 30.77 | 4 | 13 |
| 81 | 46 | 30.77 | 4 | 13 |
| 82 | 47 | 30.77 | 4 | 13 |
| 83 | 54 | 30.77 | 4 | 13 |
| 84 | 56 | 30.77 | 4 | 13 |
| 85 | 66 | 30.77 | 4 | 13 |
| 86 | 73 | 30.77 | 4 | 13 |
| 87 | 75 | 30.77 | 4 | 13 |
| 88 | 77 | 30.77 | 4 | 13 |
| 89 | 79 | 30.77 | 4 | 13 |
| 90 | 81 | 30.77 | 4 | 13 |
| 91 | 82 | 30.77 | 4 | 13 |
| 92 | 83 | 30.77 | 4 | 13 |
| 93 | 86 | 30.77 | 4 | 13 |
| 94 | 87 | 30.77 | 4 | 13 |
| 95 | 90 | 30.77 | 4 | 13 |
| 96 | 94 | 30.77 | 4 | 13 |
| 97 | 100 | 30.77 | 4 | 13 |
| 98 | 41 | 23.08 | 3 | 13 |
| 99 | 64 | 23.08 | 3 | 13 |

| | | | | |
|---|---|---|---|---|
| 100 | 65 | 23.08 | 3 | 13 |

**Table 3 List of designed mini-proteins by BindCraft**

| Design | Sequence | pLDDT | i_pAE (Å) | dG (kcal/mol) | dSASA (Å²) | Shape Complementarity | Observed m/z |
|---|---|---|---|---|---|---|---|
| BindCraft1 | TELEELFWSLSEEERPVVLR QMMERYRELERTDPELCER LDHGTTEEAAAFWTEVAR EIAAGL | 0.96 | 0.14 | -54.94 | 1624.61 | 0.72 | +5 1494.8 +6 1246.1 +7 1068.1 |
| BindCraft2 | TELEELFWSLSEEERPIVLR KMMERYRELERTDPCLAE RLDHGTTEEAAAFWTEVA REIHESL | 0.95 | 0.14 | -58.38 | 1649.82 | 0.71 | +6 1264.1 +7 1083.6 +8 948.3 |
| BindCraft3 | MKEIAEKVWLVSQYILENP EDEELWKECHEAQKHPEE MMKFVEEKAEEIEKEK | 0.94 | 0.17 | -55.57 | 1476.24 | 0.74 | +5 1305.0 +6 1087.7 +7 932.6 |
| BindCraft4 | MEEIAEKVWLVSQYILENP EDEELWKKCHEAQKHPED MMKFVEEEAEKLKKKE | 0.93 | 0.17 | -54.23 | 1530.72 | 0.74 | +5 1301.8 +6 1085.0 +7 930.2 |
| BindCraft5 | SFEHVKKELEKELSKNDKH QEPYIPIGIEMFKTMIENPEY SHGFTPEEFEELIELLEK | 0.92 | 0.17 | -60.66 | 1917.38 | 0.67 | +5 1396.3 +6 1163.7 +7 997.5 |
| BindCraft6 | SFEHVLEELKKELAKNDKH QEPYIPIGIEMFKTMICNPEY SHGFTPEQFEELIKLLQE | 0.92 | 0.17 | -63.16 | 1978.81 | 0.63 | +5 1384.7 +6 1154.0 +7 989.3 |
| BindCraft7 | DPSPEKYREIYPELRSHQEE YDKSPECQEEVVAKLKEFL KEKGVLK | 0.94 | 0.16 | -56.54 | 1554.06 | 0.71 | +5 1105.4 +6 921.4 +7 790.2 |
| BindCraft8 | TEEDRMNHKIATEIDWDWF VKFHKEMEEYRKRHPNCS DKELALYGMKKIIEYFEKN VKDEEVLKYLEKMKKSTE | 0.94 | 0.16 | -70.75 | 2012.21 | 0.69 | +7 1319.5 +8 1154.6 +9 1026.6 |
| BindCraft9 | SKMEKEAEEVKEKLLEYIK DMPEEERKANEECVEAHIK KIIDIYESGKEEEARMWIDL YKKMF | 0.94 | 0.16 | -53.81 | 1599.34 | 0.74 | +6 1281.2 +7 1098.3 +8 961.2 |
| BindCraft10 | DPSPEKYREIYPELRSHQAE YDKSPECQKQVVEKLKKY LKEKGVLK | 0.94 | 0.16 | -56.39 | 1552.72 | 0.74 | +5 1108.2 +6 923.7 +7 791.9 |

Note: Residues in red are A to C mutations to allow mini-protein synthesis via native chemical ligation.

**Table 4 List of designed mini-proteins by Odesign**

| Design | Sequence | Binder PTM | Min iPAE PTM (Å) | ddG (kcal/mol) | Contact Molecular Surface (Å²) | Observed m/z |
|---|---|---|---|---|---|---|
| Odesign1 | EELKKELERLEERLKRLEEALACETD PEEKASIQAAIEETKKKIEEVKAKL | 0.82 | 1.11 | -32.23 | 305.83 | +5 1194.2 +6 995.4 +7 853.3 |
| Odesign2 | MTVEELEAQMEPVAARIYAIARRDP AAWARVRAACQQFPPIPENPLPEVL EQVLAQLRALEAELA | 0.84 | 0.94 | -57.24 | 503.49 | +5 1451.9 +6 1210.0 +7 1037.1 |

| | | | | | |
|---|---|---|---|---|---|
| Odesign3 | MTVEEVLAEMEPVARAVFAICRRDP<br>AAWARVRAAACTFPPIPETPEVETLE<br>AVLAGLRALLAELA | 0.82 | 0.94 | -53.8 | 527.05 | +5 1395.2<br>+6 1162.6<br>+7 996.8 |
| Odesign4 | SVTAAELAALVAAARRAQAALLAA<br>GEVEPMPDAELRAAVAATLAALAA<br>RGVRPADVIAEIEAAIA | 0.72 | 1.51 | -54.75 | 452.41 | NA |
| Odesign5 | SALQAQIERLEAELAALKQALAEET<br>DPEEKKSIQLAIEKTEEELARLKAEA | 0.77 | 1.4 | -27.78 | 272.5 | NA |
| Odesign6 | SELQQQIERLEAELAALRKAYEEETD<br>PEEKASIEKAIEETEKKLEELKKKL | 0.78 | 1.32 | -25.13 | 241.92 | NA |

Note: Residues in red are A to C mutations to allow mini-protein synthesis via native chemical ligation.

### HPLC and MST analysis of BindCraft1

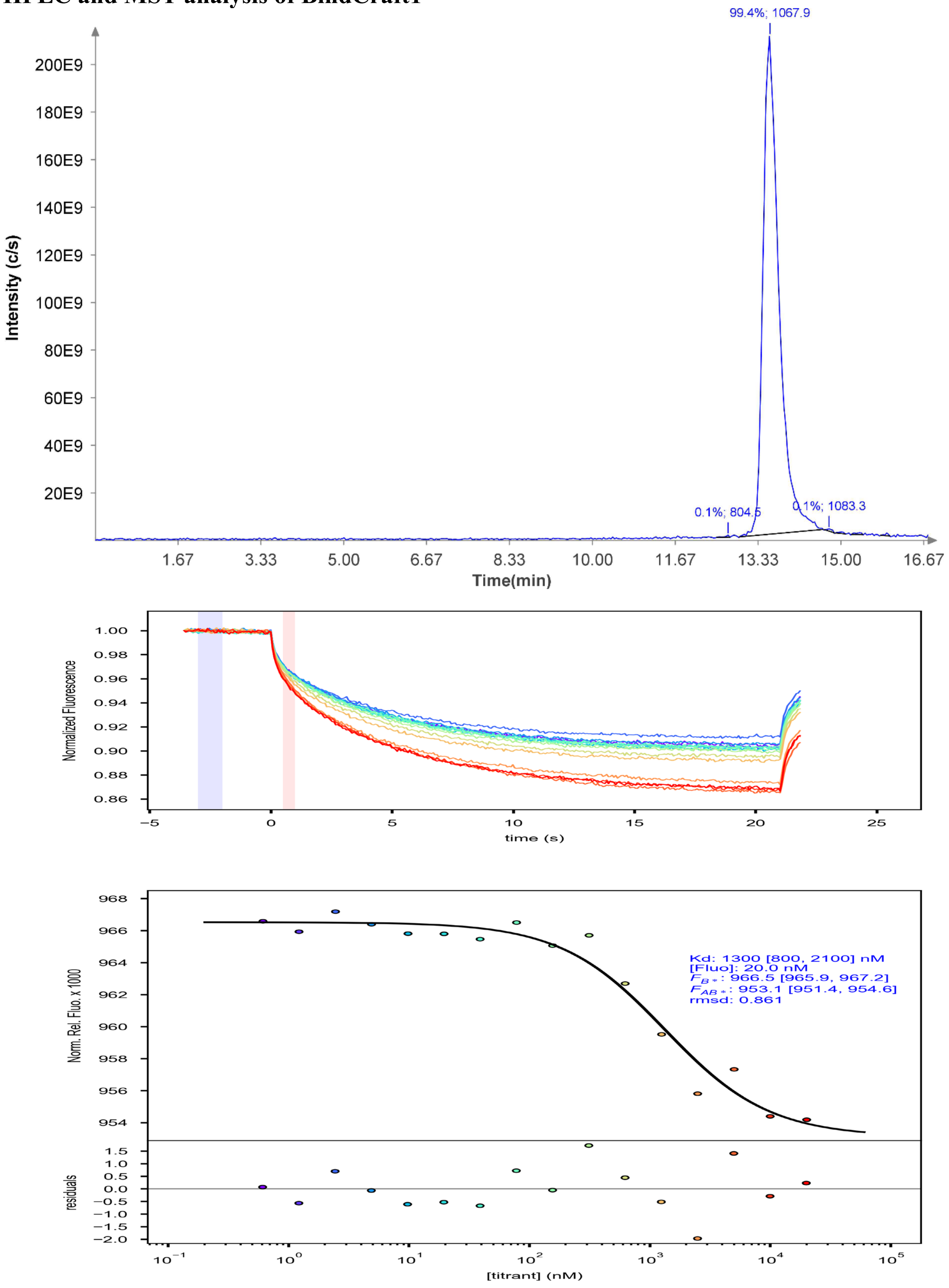

#### HPLC and MST analysis of BindCraft2

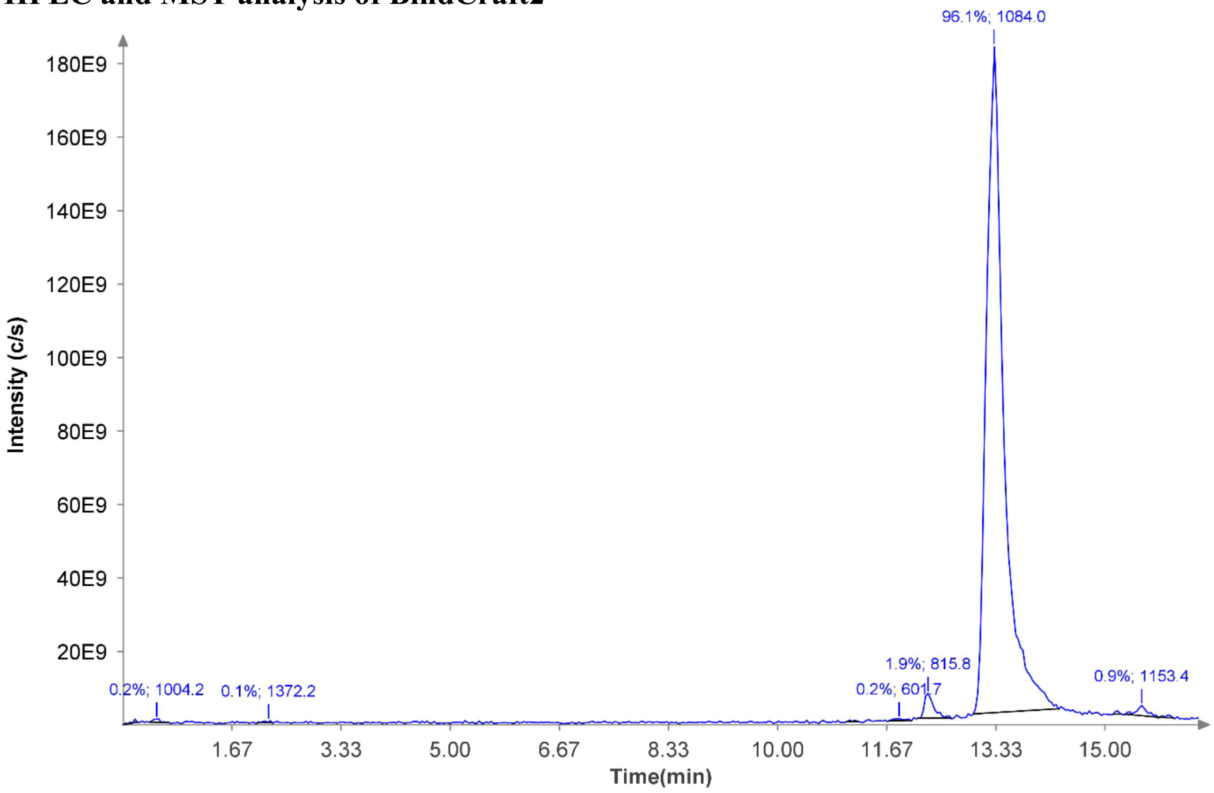


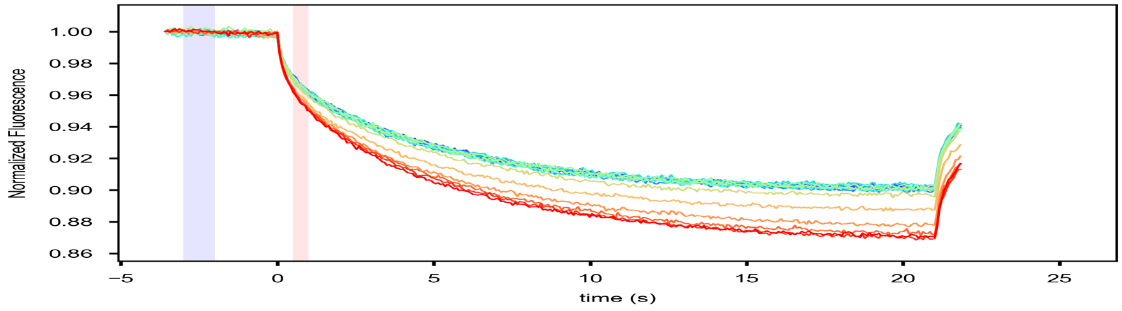


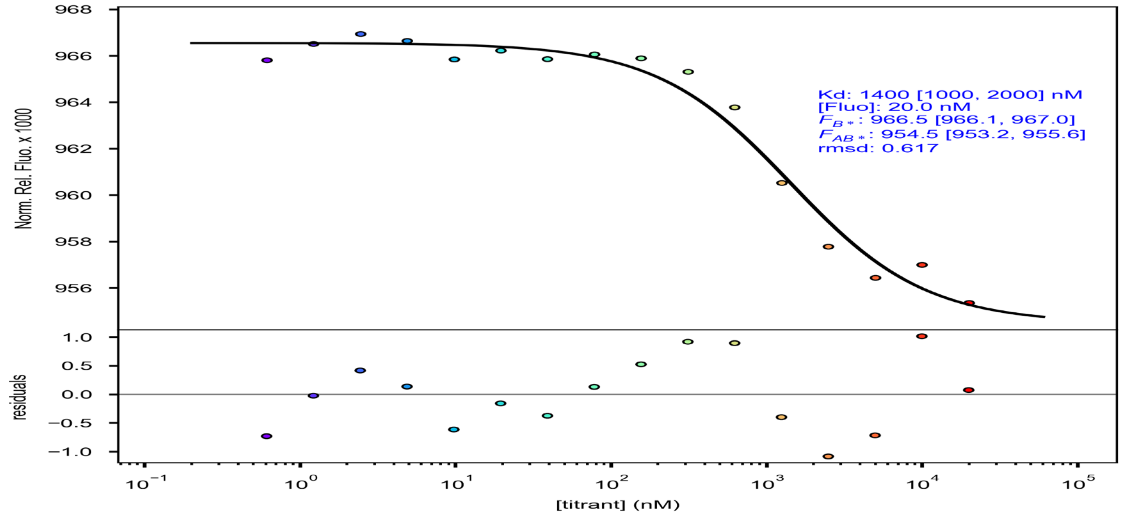

#### HPLC and MST analysis of BindCraft3

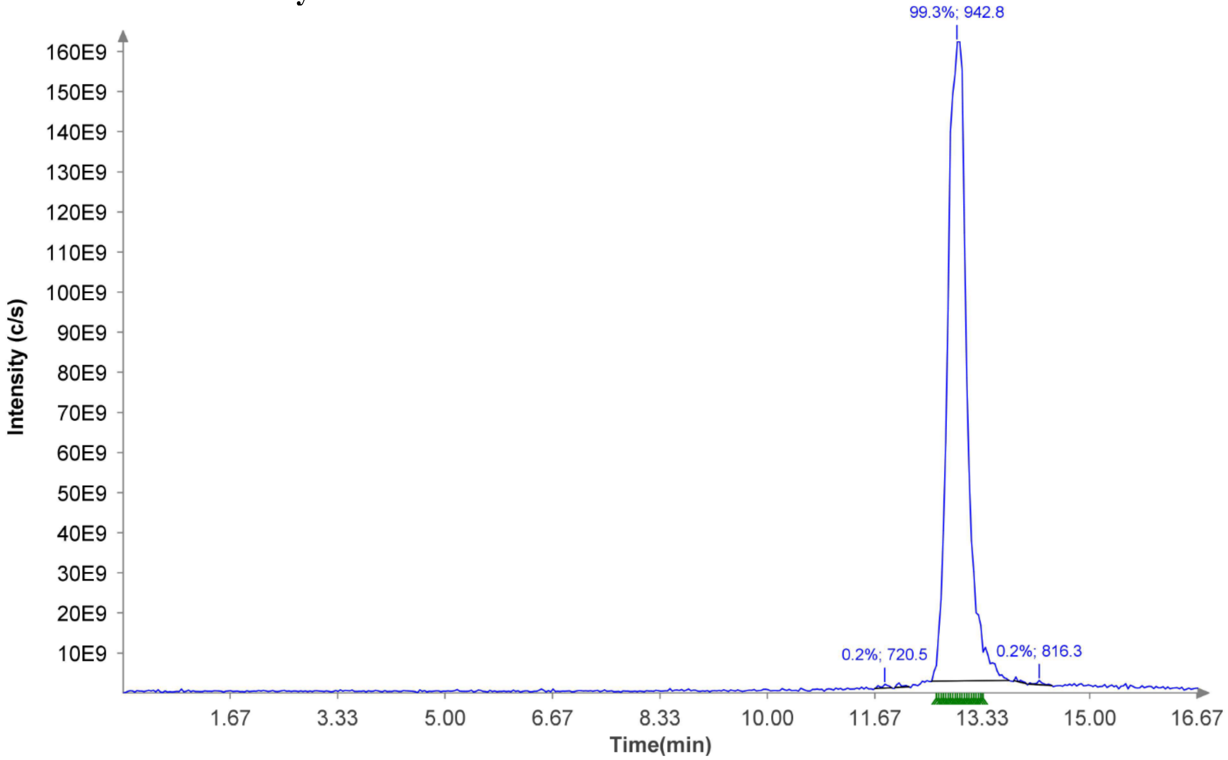


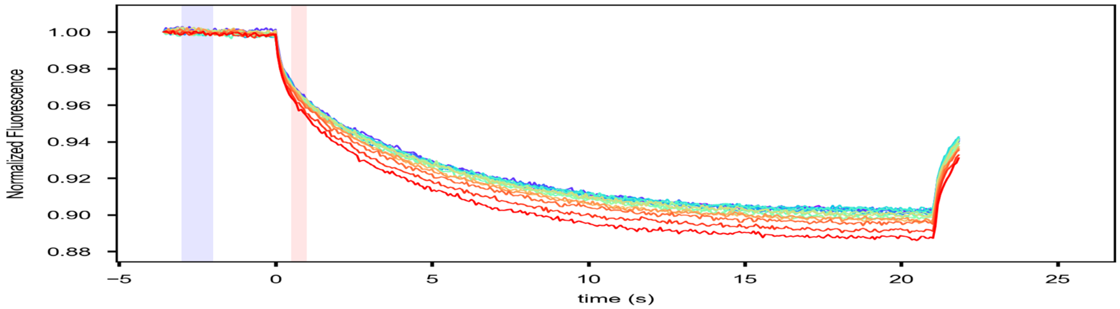


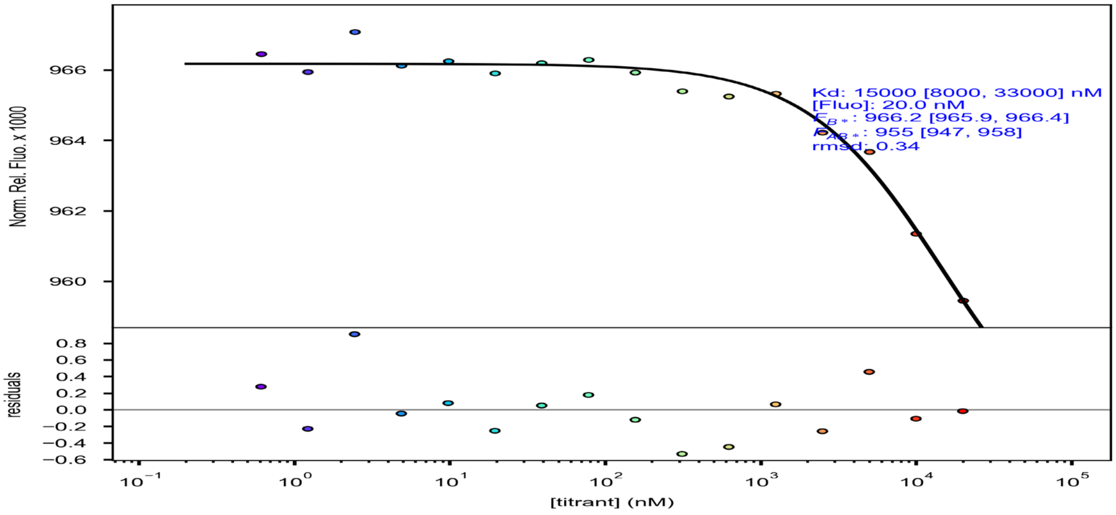

### HPLC and MST analysis of BindCraft4

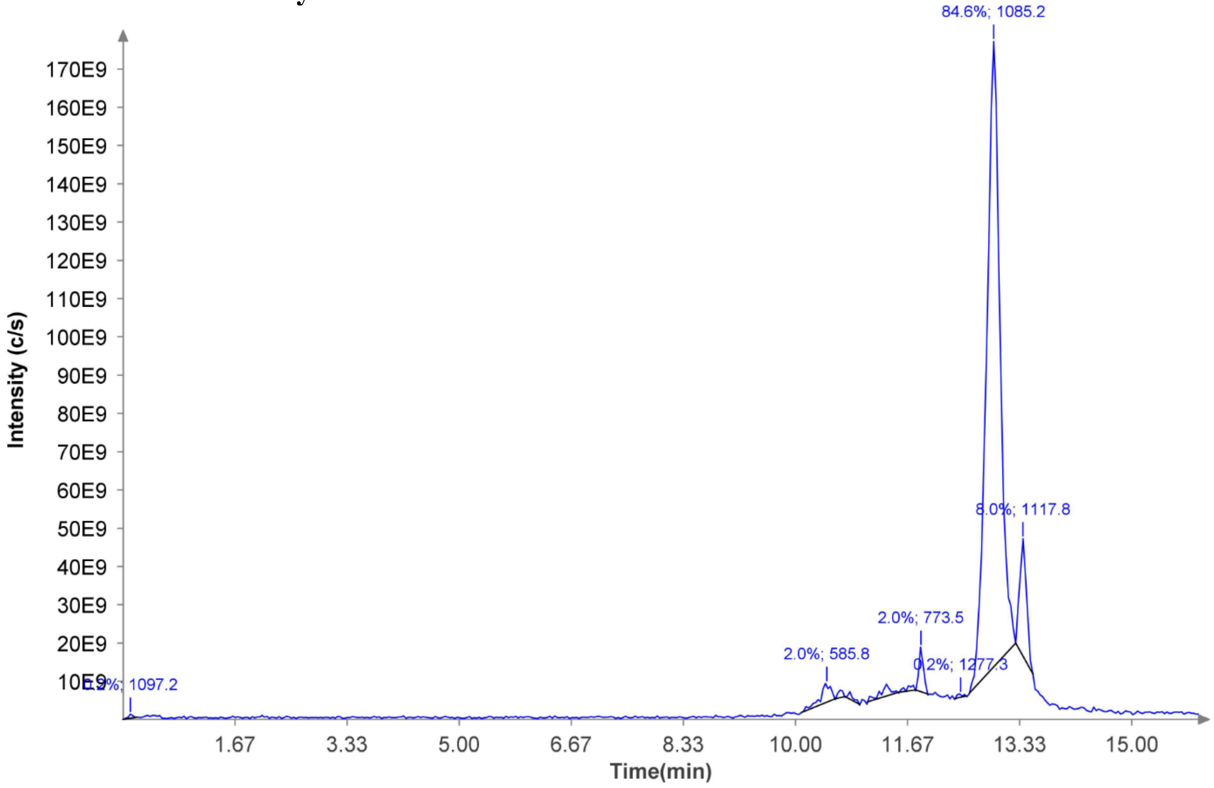


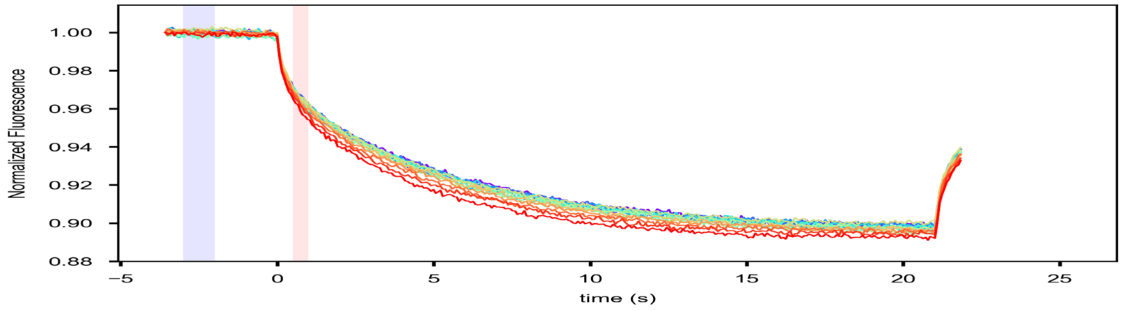


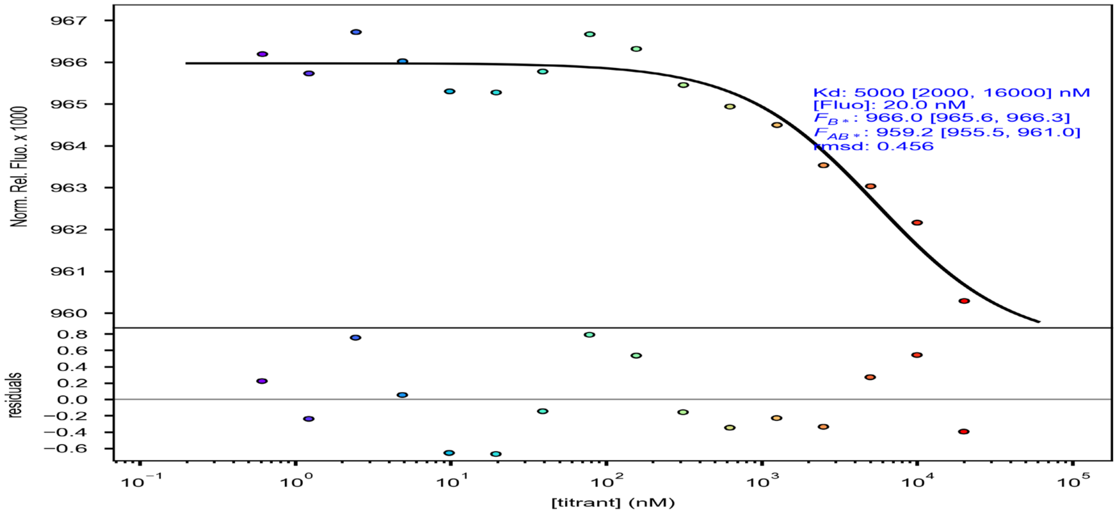

### HPLC and MST analysis of BindCraft5

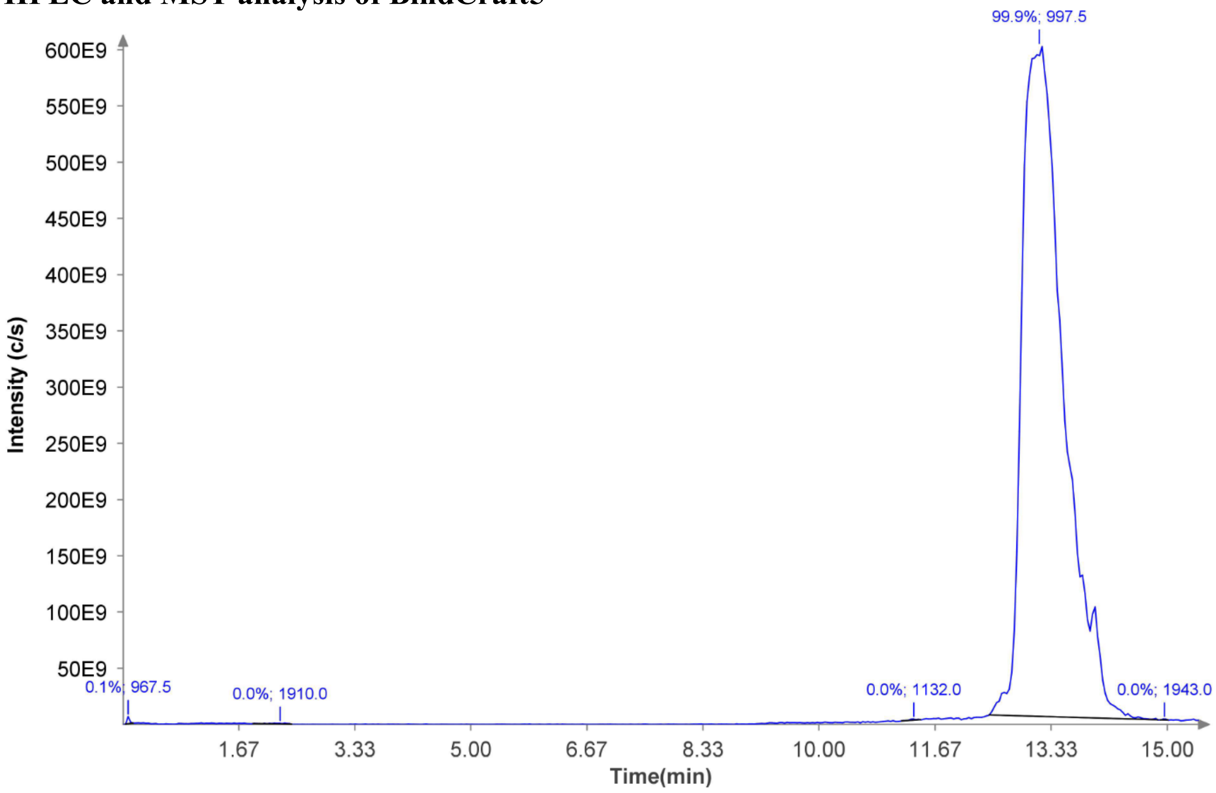


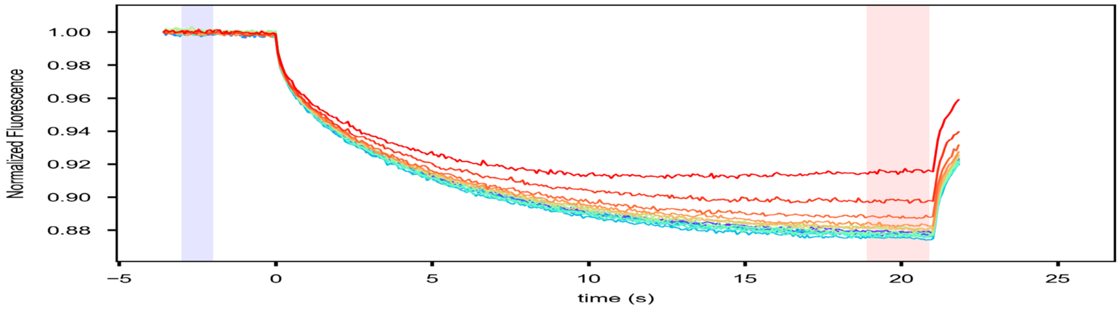


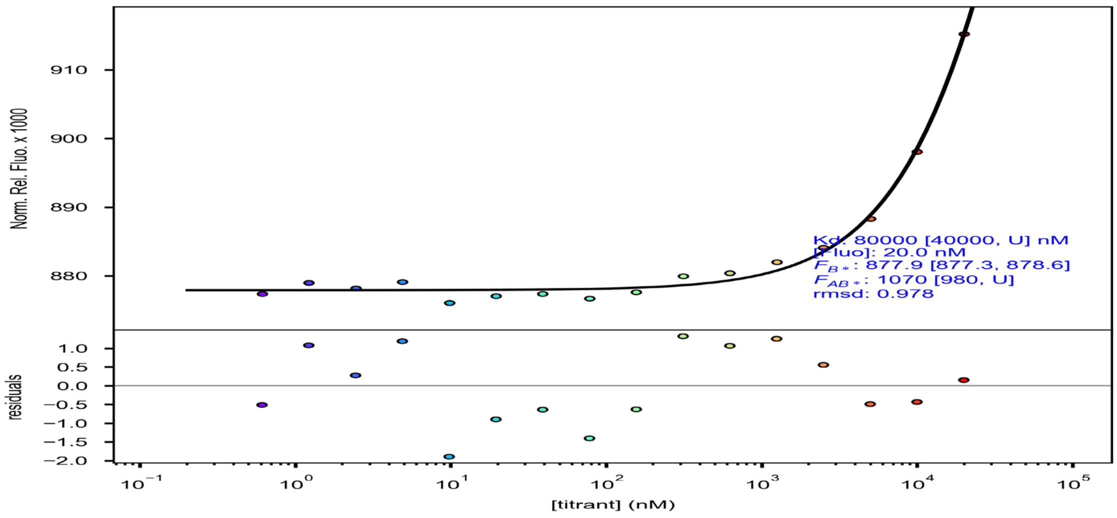

### HPLC and MST analysis of BindCraft6

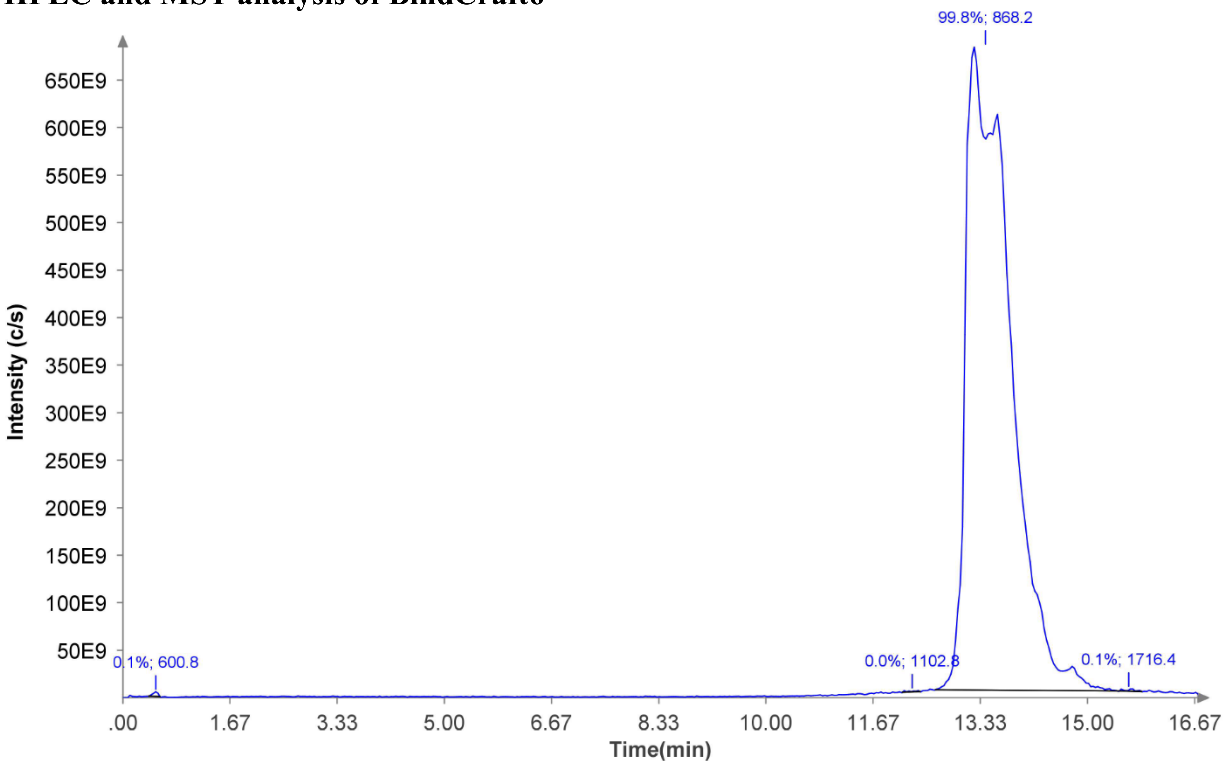


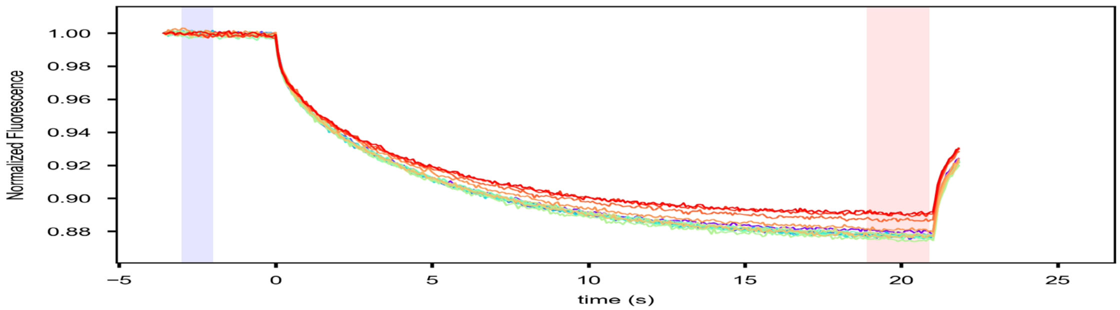


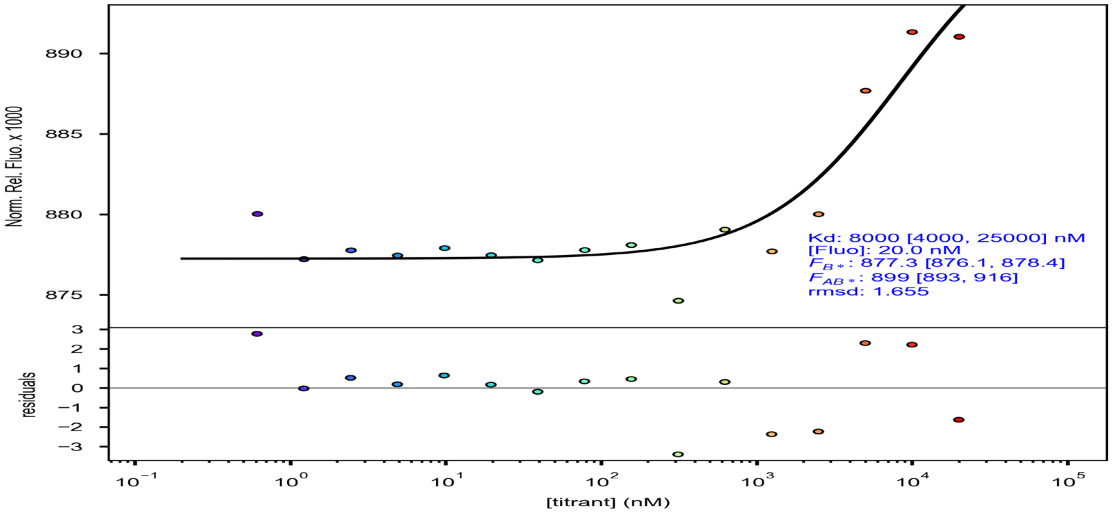

#### HPLC and MST analysis of BindCraft7

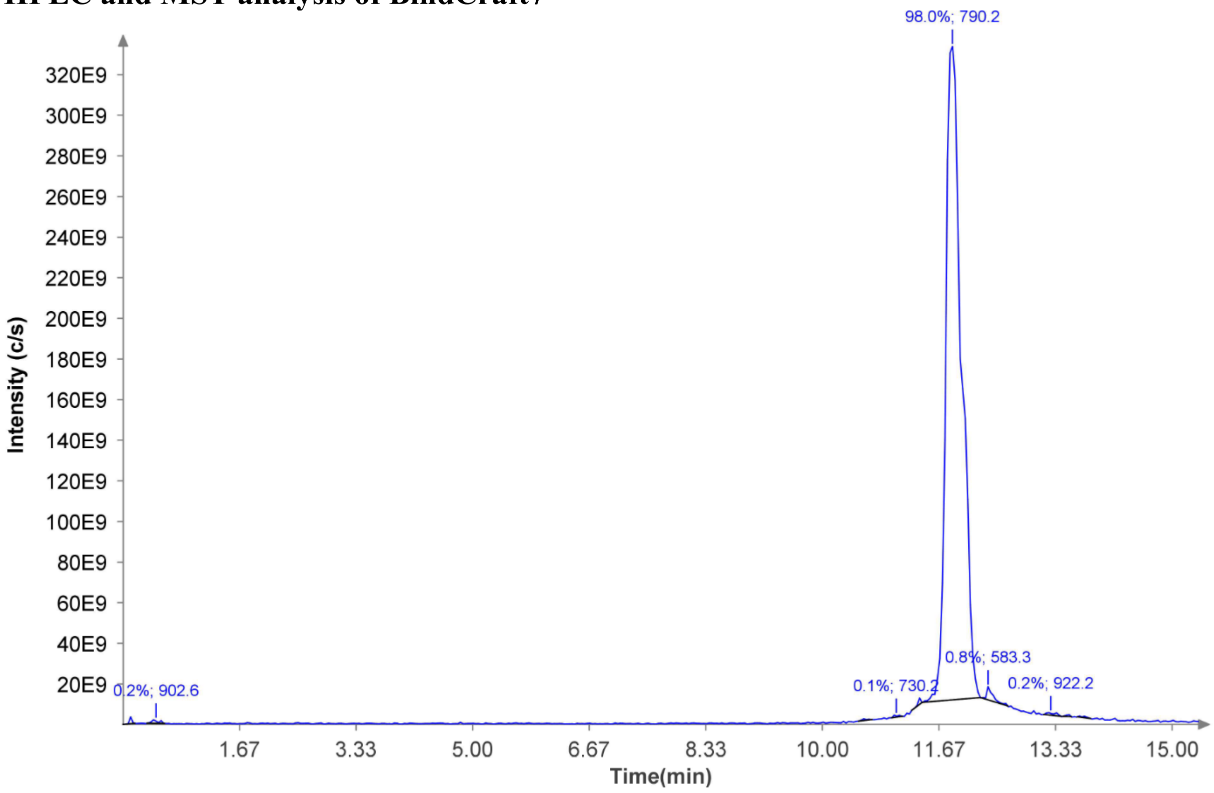


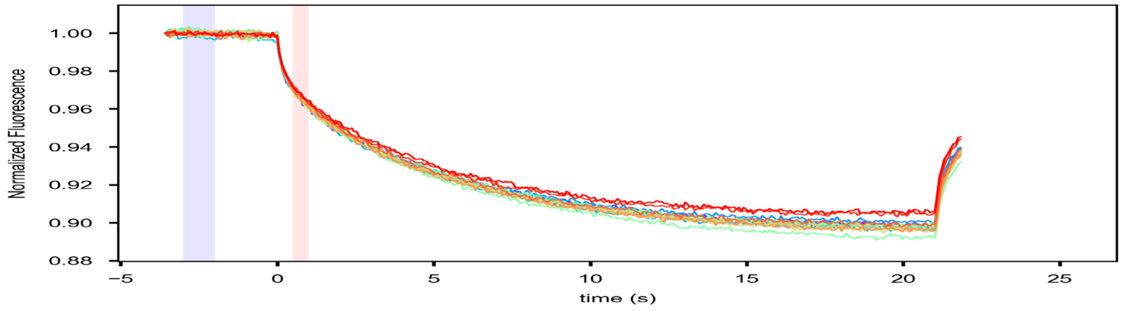


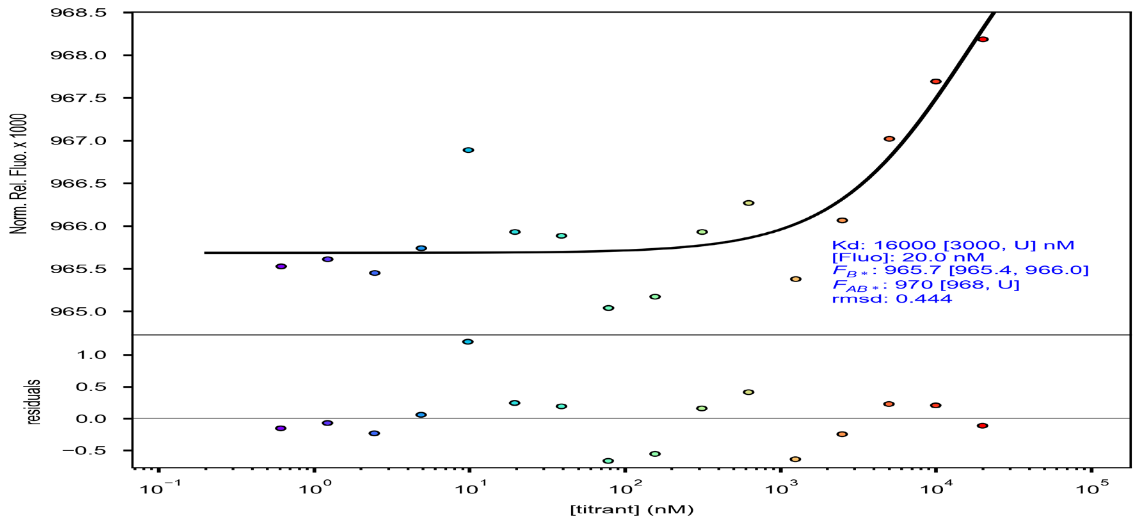

### HPLC and MST analysis of BindCraft8

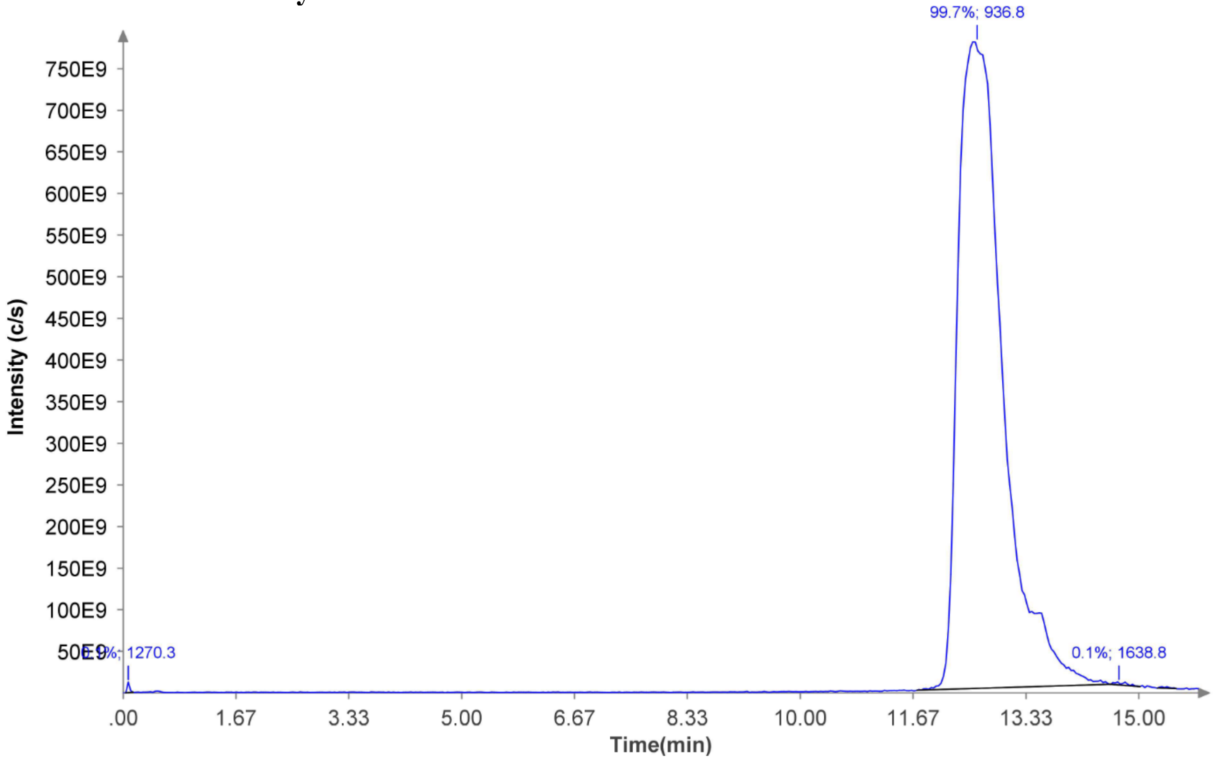


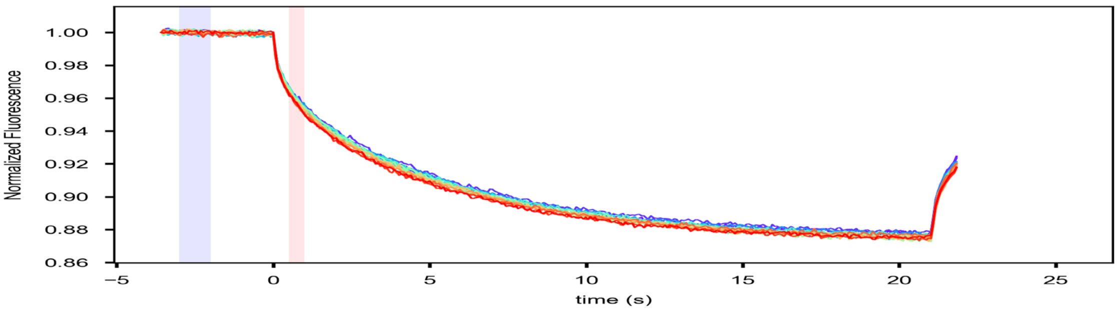


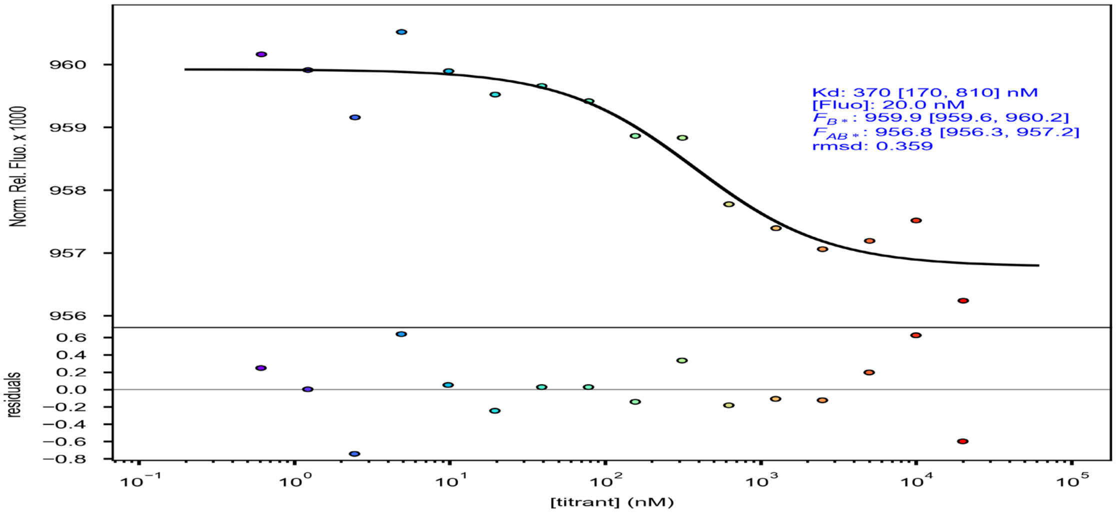

**HPLC and MST analysis of BindCraft9**

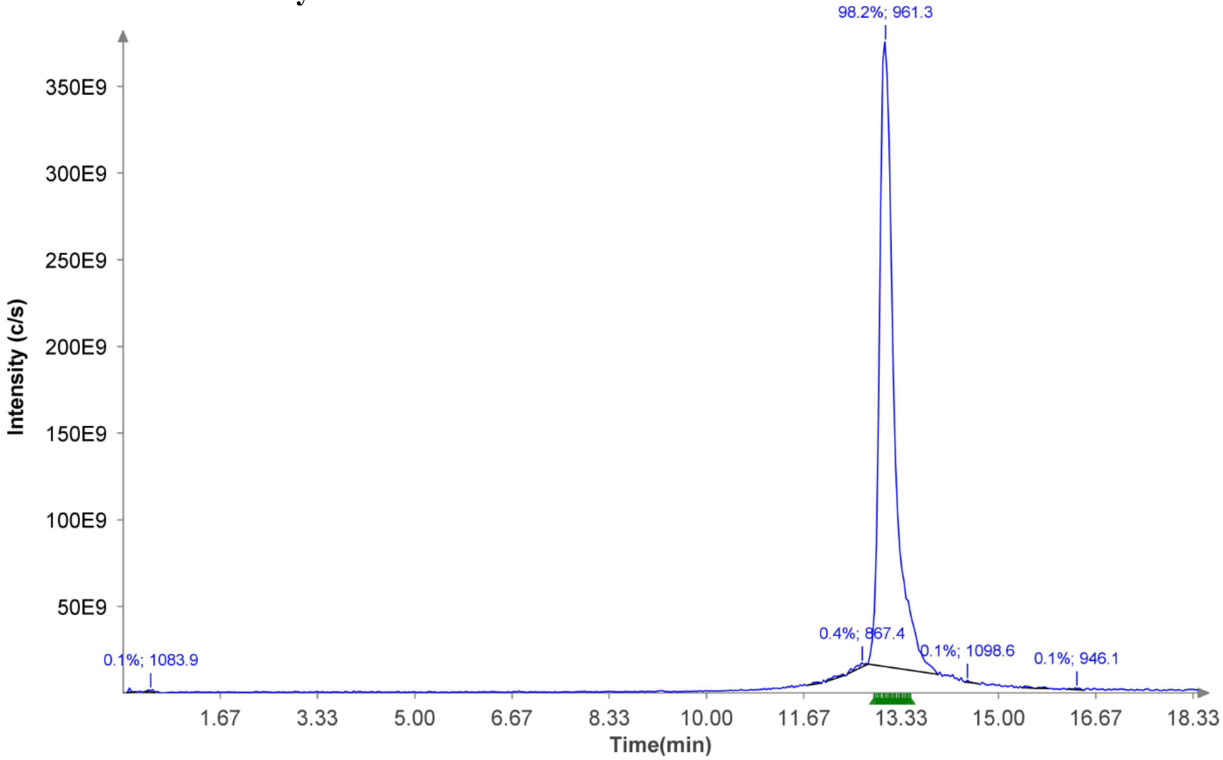


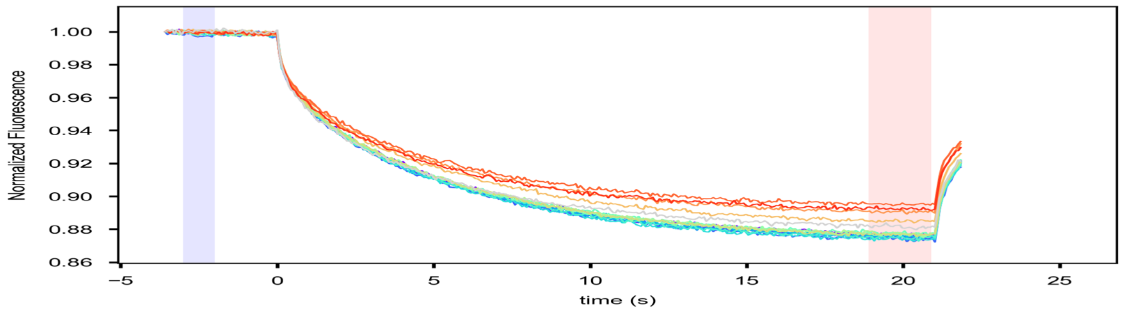


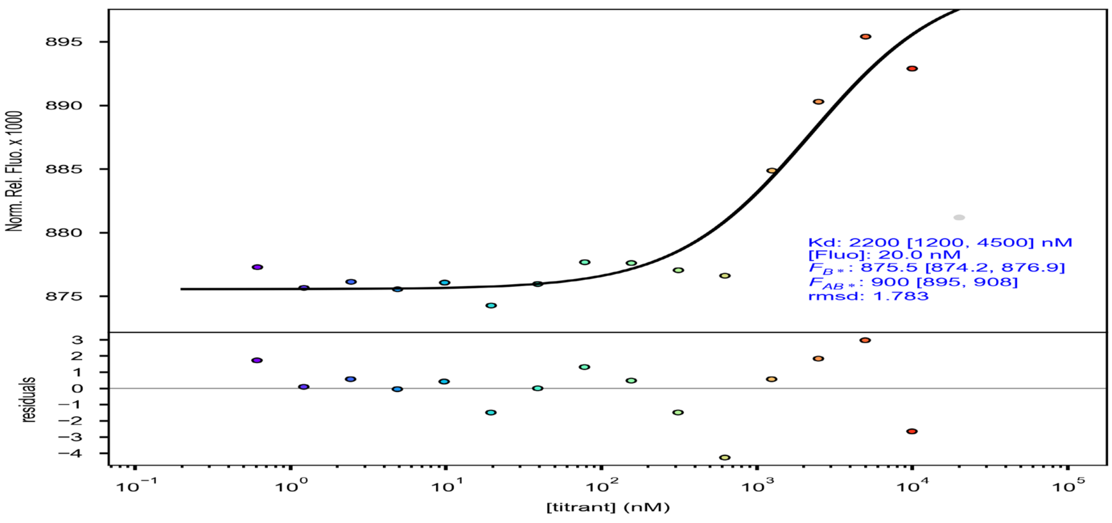

#### HPLC and MST analysis of BindCraft10

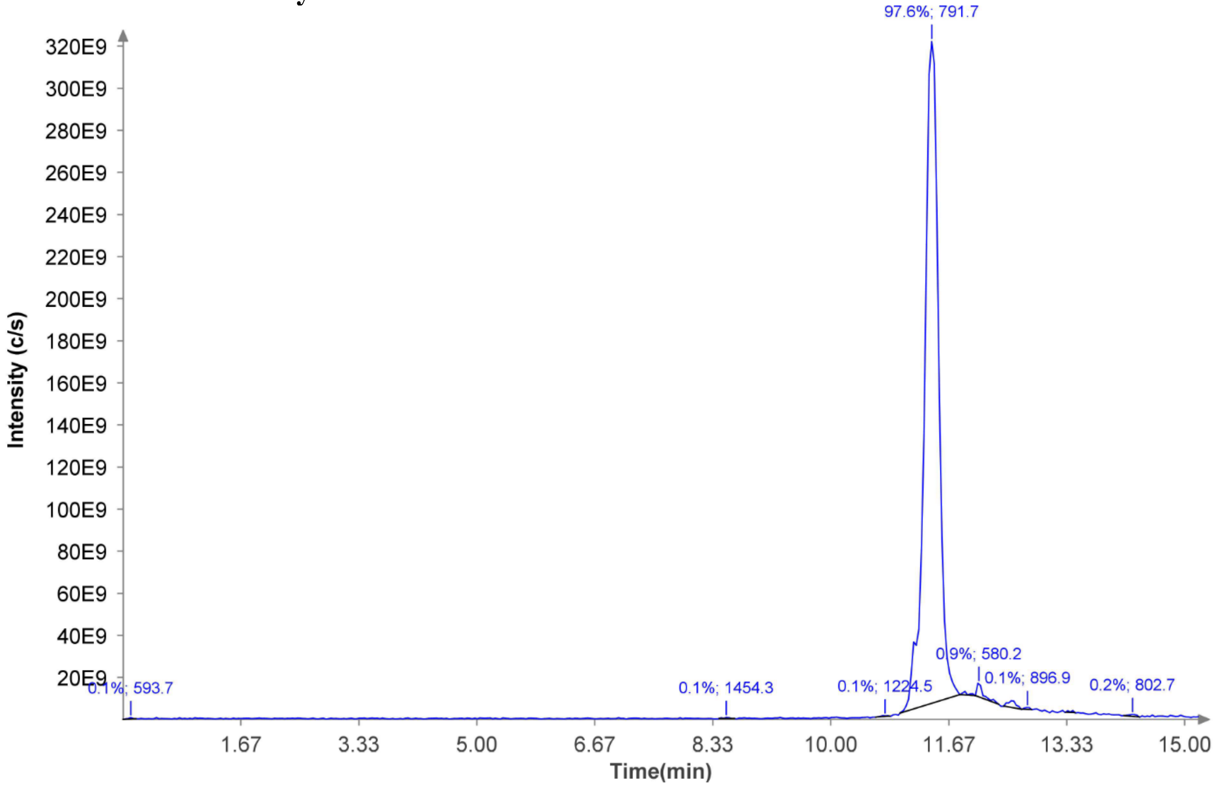


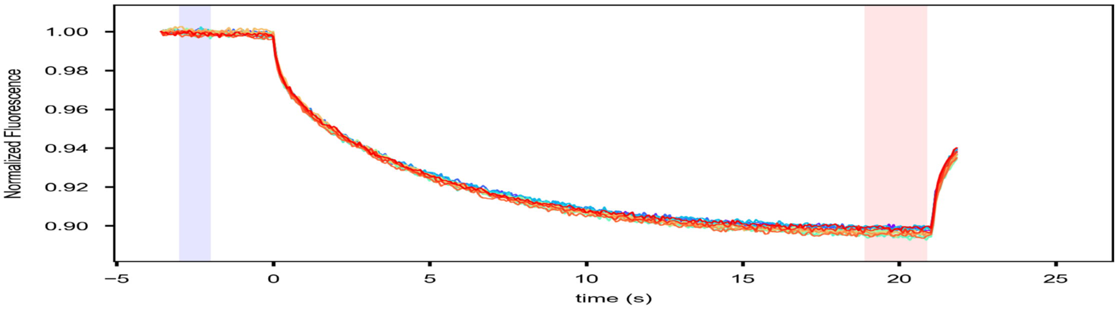


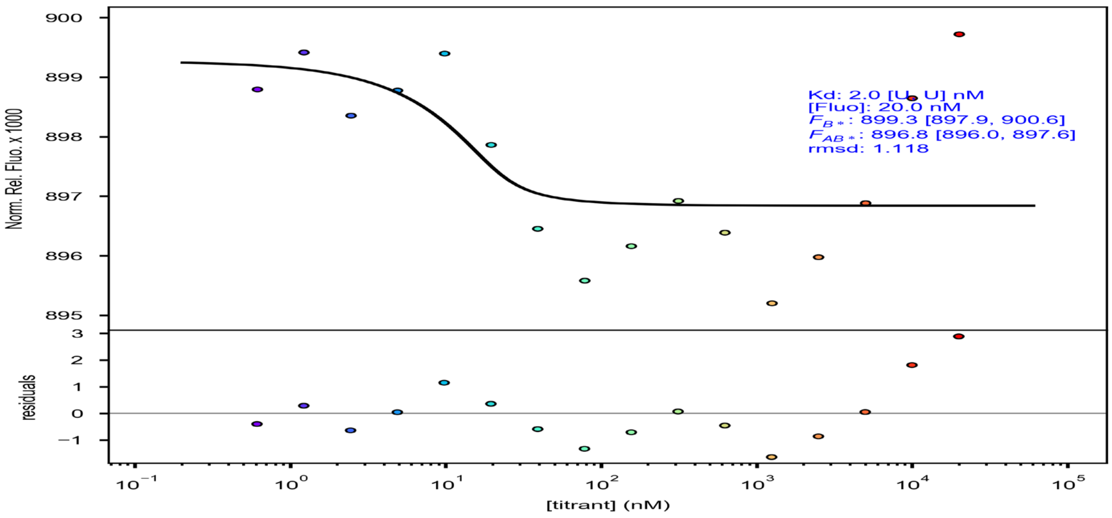

#### HPLC and MST analysis of Odesign1

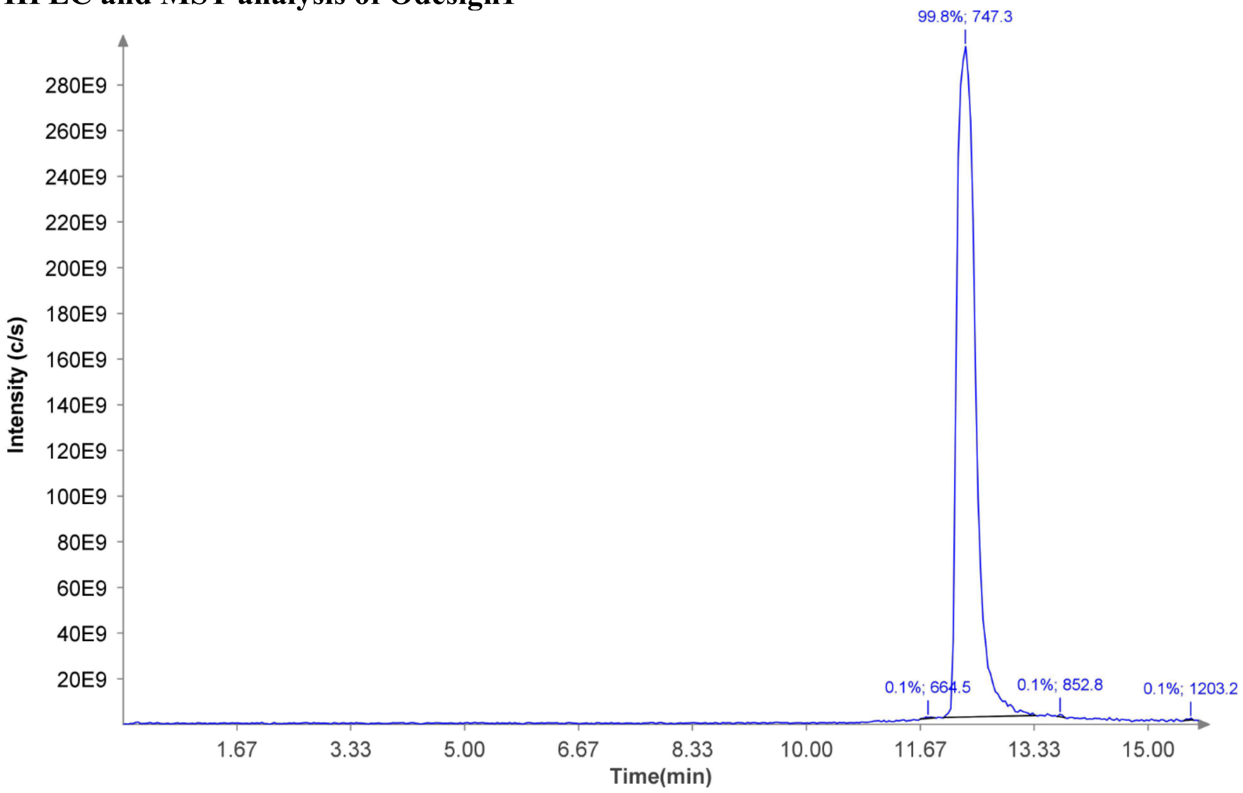


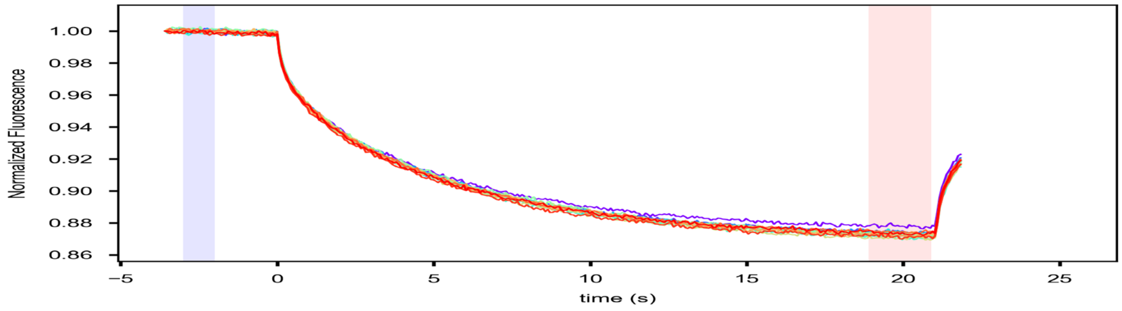


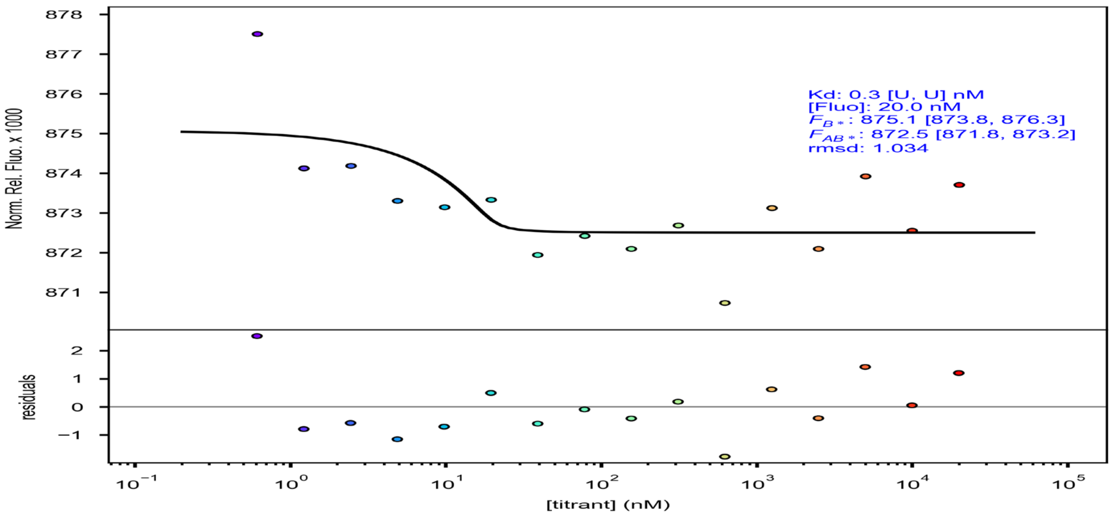

### HPLC and MST analysis of Odesign2

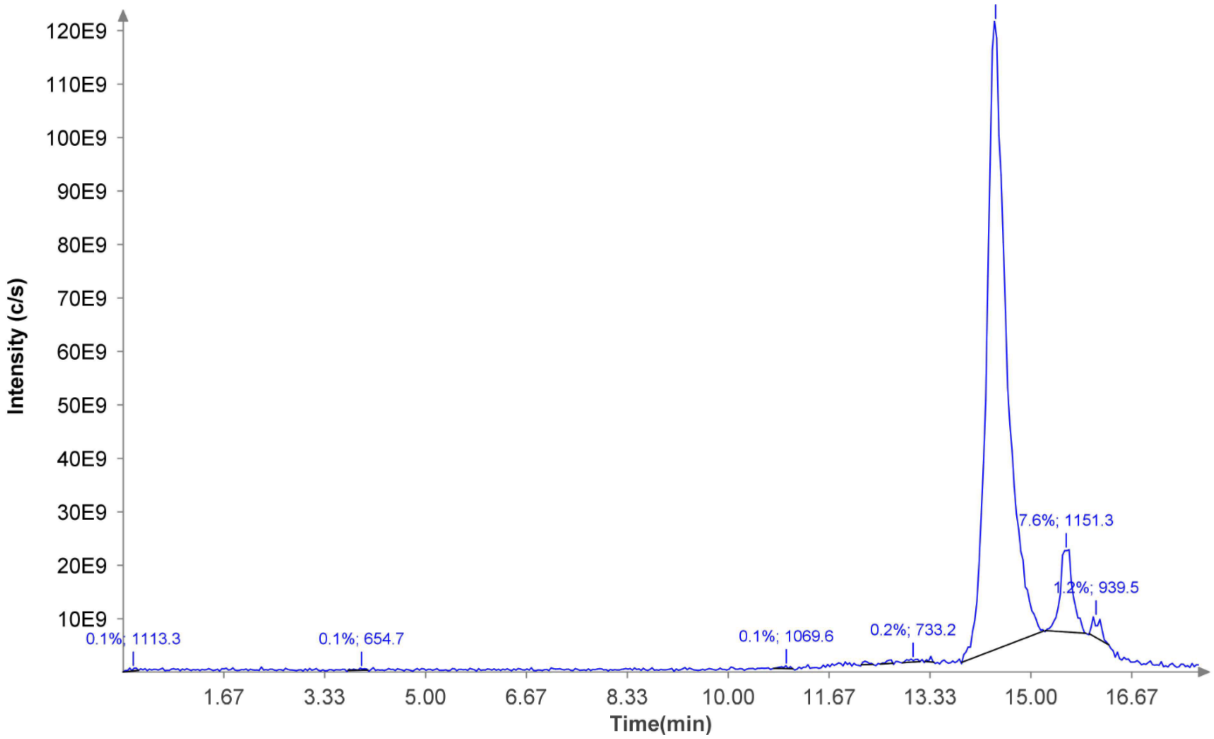


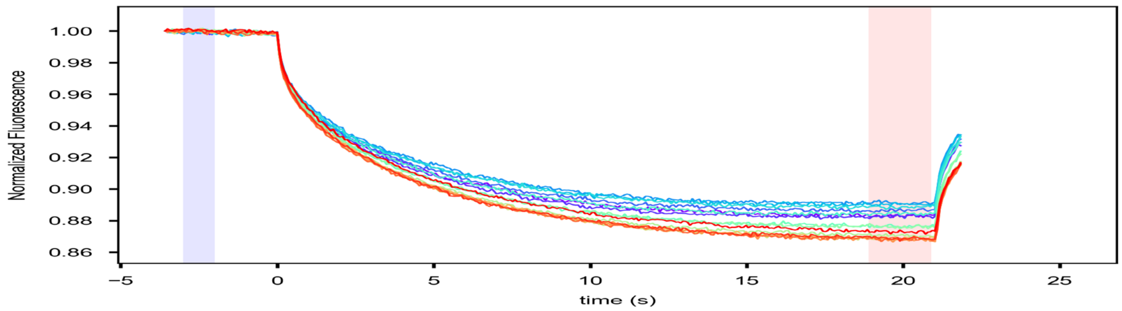


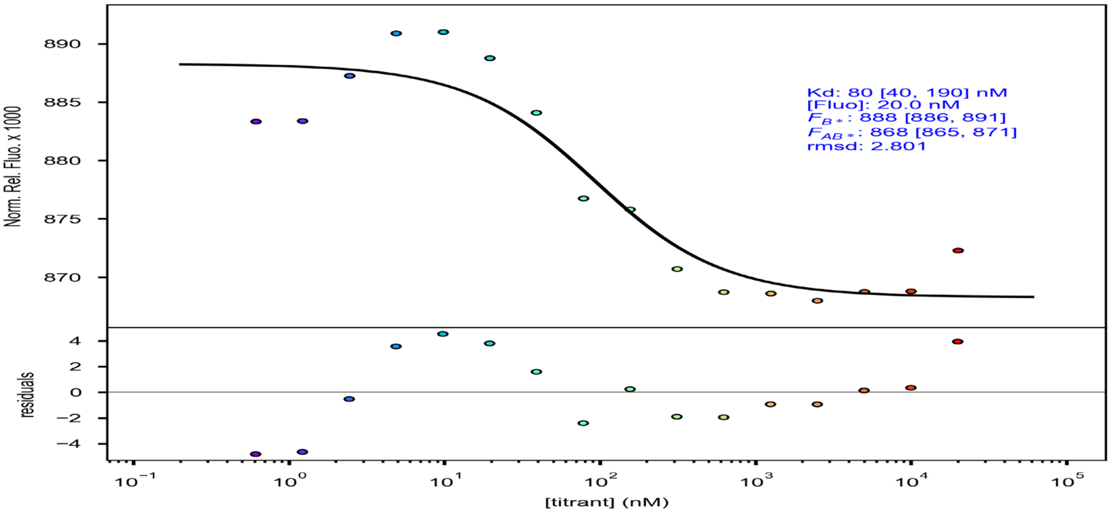

#### HPLC and MST analysis of Odesign3

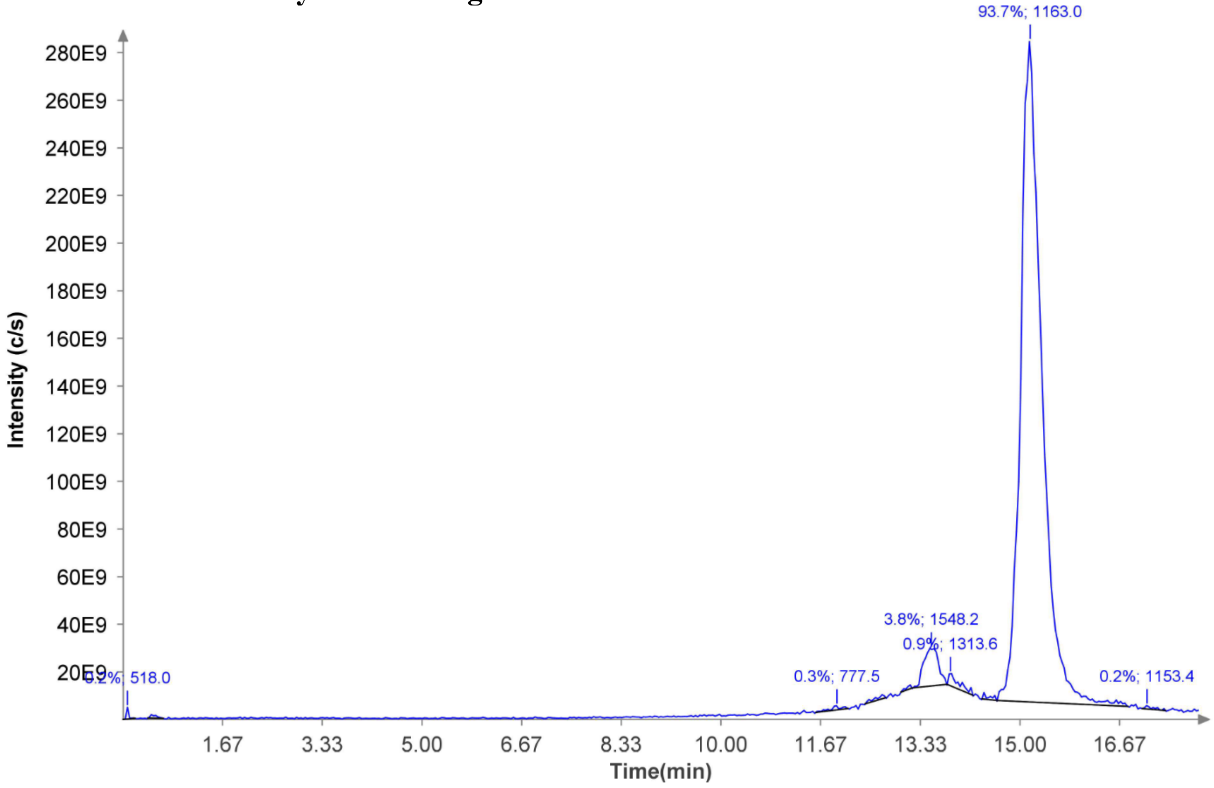


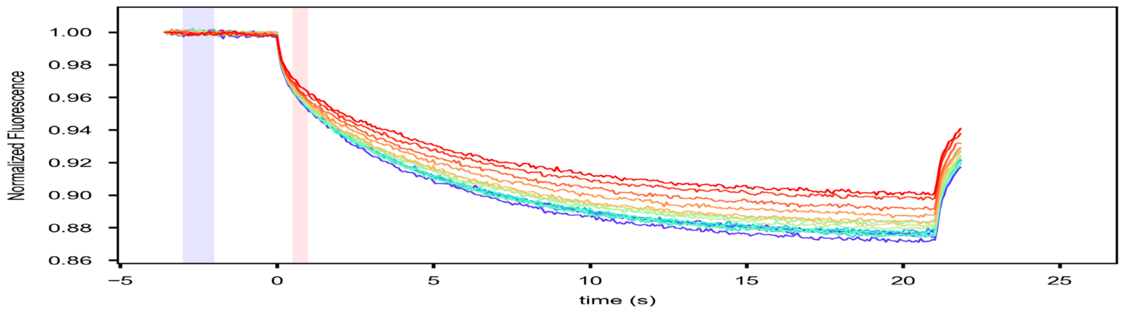


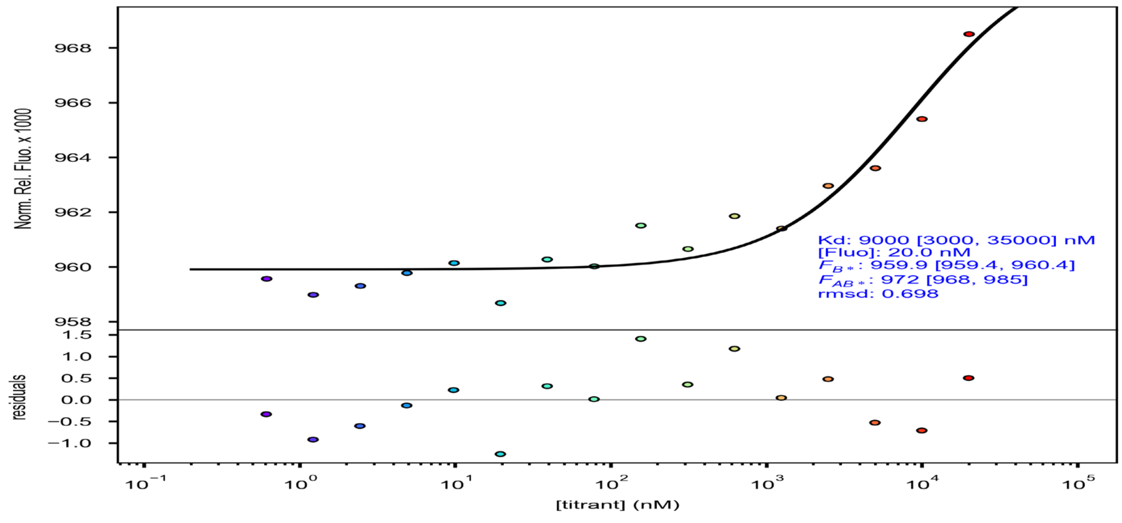
